# Distinct pathogenic genes causing intellectual disability and autism exhibit overlapping effects on neuronal network development

**DOI:** 10.1101/408252

**Authors:** Monica Frega, Martijn Selten, Britt Mossink, Jason M. Keller, Katrin Linda, Rebecca Moerschen, Jieqiong Qu, Pierre Koerner, Sophie Jansen, Elske Bijvank, Astrid Oudakker, Tjitske Kleefstra, Hans van Bokhoven, Huiqing Zhou, Dirk Schubert, Nael Nadif Kasri

## Abstract

An intriguing question in medical biology is how mutations in functionally distinct genes can lead to similar clinical phenotypes. For example, patients with mutations in distinct epigenetic regulators EHMT1, MBD5, MLL3 or SMARCB1 share the core clinical features of intellectual disability (ID), autism spectrum disorder (ASD) and facial dysmorphisms. To elucidate how these phenotypic similarities are reflected by convergence at the molecular, cellular and neuronal network level, we directly compared the effects of their loss of function in neurons. Interestingly, knockdown of each gene resulted in hyperactive neuronal networks with altered patterns of synchronized activity. At the single-cell level, we found genotype-specific changes in intrinsic excitability and excitatory-inhibitory balance, but in all cases leading to increased excitability. Congruent with our physiological findings, we identified dysregulated genes that converge on biological and cellular pathways related to neuronal excitability and synaptic function, including genes previously implicated in ID/ASD. Yet, our data suggests that the common cellular phenotypes depend on the ensemble of dysregulated genes engaged in neuronal excitability rather than the direction of transcriptional changes of individual genes. The demonstration of increasing convergence from molecular pathways to neuronal networks may be a paradigm for other types of ID/ASD.

## Introduction

Neurodevelopmental disorders (NDDs), including intellectual disability (ID) and autism spectrum disorder (ASD), are genetically and phenotypically heterogeneous. Despite the identification of Mendelian mutations in more than 800 genes that give rise to some type of NDD(1), our understanding of the key molecular players and mechanisms is still fragmented and needs conceptual advances. Furthermore, how mutations and DNA variants in distinct genes can, in some cases, lead to similar clinical phenotypes, is poorly understood(2, 3). Recent studies have proposed that the genetic heterogeneity among NDDs is buffered at the level of molecular pathways where the effects of many different DNA variants converge(4–6). However, we still have to resolve the exact nature of such converging pathways and how disruptions thereof give rise to commonality in terms of brain dysfunction and pathology.

In recent years, evidence has accumulated that synaptic processes and neuronal gene transcription through epigenetic modification of chromatin structure plays an important role in both normal cognitive processes and the etiology of NDDs(3, 7). Kleefstra syndrome (OMIM#610253) is an example of a rare NDD comprising ID, ASD, hypotonia and dysmorphic features as major hallmark phenotypes(8–10). The canonical disease is caused by *de novo* loss-of-function mutations in the gene *EHMT1* (Euchromatin Histone Lysine Methyltransferase 1, also known as GLP)(8). Interestingly, however, we previously found that *de novo* mutations in four other chromatin modifiers, i.e. *SMARCB1* (SWI/SNF-related matrix-associated actin-dependent regulator of chromatin, subfamily B member 1), *MLL3* (Histone-lysine N-methyltransferase 2C, or *KMT2C), NR1I3* (Nuclear receptor, subfamily 1 group I member 3) and *MBD5* (Methyl-CpG-binding domain protein 5), result in core clinical features highly reminiscent of Kleefstra syndrome and that we collectively refer to as the Kleefstra syndrome phenotypic spectrum (KSS)(11). The corresponding proteins are directly or indirectly involved in epigenetic regulation of gene expression and are also associated with other disorders that share certain cognitive features with KSS. For example, *MBD5* deletions are associated with the chromosome 2q23.1 deletion syndrome resembling Smith-Magenis syndrome(12, 13), missense mutations in *SMARCB1* are associated with Coffin-Siris syndrome(14, 15), and intragenic *EHMT1* duplications are associated with schizophrenia(16, 17).

EHMT1 cooperates with its mammalian paralogue EHMT2/G9a and exhibits enzymatic activity for histone 3 lysine 9 mono and di-methylation (H3K9me1 and H3K9me2, respectively), which is known to promote a heterochromatic structure and hence gene repression(18, 19). Loss of EHMT1 function in mice and *Drosophila* leads to learning and memory impairments(20–24). Additionally, *Ehmt1^+^* mice recapitulate autistic-like features that are seen in patients with Kleefstra syndrome(20). At the cellular level, these mice show a significant reduction in dendritic arborisation and the number of mature spines in CA1 pyramidal neurons(21), together with a reduced ability to establish synaptic scaling, a specific form of homeostatic plasticity(25). Furthermore, EHMT1 deficiency alters cortical neuronal network activity during development(26), but the underlying mechanisms remain to be determined.

Each of the KSS gene products functions to epigenetically regulate transcription, while protein-protein interaction data and genetic interaction studies in *Drosophila* indicate that the corresponding proteins are engaged in shared biological processes(11). A recent study in *Drosophila* has strengthened this notion by showing that two of the KSS genes, *EHMT1* and *KMT2C (MLL3)*, are required for short-term memory and share direct and indirect gene targets(27). Collectively this leads to the hypothesis that the epigenetic modifiers associated with KSS coalesce on gene networks for molecular or cellular pathways that affect neuronal function in the same way. Yet, this hypothesis is seemingly confounded by the fact that the modifiers have distinct and in some cases even antagonistic functions(27). For example, EHMT1 and MLL3 directly modify histones(28). But the H3K9me1 and H3K9me2 marks catalyzed by EHMT1 repress gene transcription, while H3K4 methylation by MLL3 results in transcriptional activation. Furthermore, SMARCB1 is part of an ATP-dependent chromatin remodeling complex(29, 30), MBD5 binds to heterochromatin(31), and NR1I3 is a nuclear hormone receptor(32).

In this study, we combined molecular, cellular and electrophysiological approaches to address the question of whether a loss in any KSS gene similarly affects neuronal function. We directly compared monogenic loss of four KSS genes (*Ehmt1, Smarcb1, Mll3* and *Mbd5)* in developing neuronal networks. We show that despite several functional and molecular changes unique to each respective KSS gene knockdown, all of the KSS gene-deficient neuronal networks were hyperactive during the course of development and showed an altered organization compared to wildtype networks. At the single-cell level, we show that cortical neurons were intrinsically more excitable and received less inhibitory input, thereby shifting the excitatory-inhibitory (E/I) balance. At the transcriptional level, KSS gene deficiency converged on biological and cellular pathways related to potassium channels, synaptic transmission and the post-synaptic density, albeit, distinguishable gene expression patterns were observed between genotypes. Finally, we confirmed that reduced inhibitory synaptic input and increased intrinsic excitability also led to hyperexcitability in *Ehmt1^+/-^* mice, a mouse model for Kleefstra syndrome(20). In the context of integrated analysis of NDDs caused by haploinsufficiency in interrelated chromatin pathways, our results may provide a first explanation for why core clinical features are shared by KSS patients and other phenotypically congruent, but genetically distinct disorders involving ID and ASD.

## Results

### Knockdown of KSS genes leads to hyperactive neuronal network activity during development

We investigated whether the loss of function of individual KSS genes (*Ehmt1, Smarcb1, Mll3* or *Mbd5*) leads to neuronal circuitry impairments *in vitro. NR1I3* was not included since we found it not to be expressed in primary rat cortical neurons (data not shown). To compare neuronal networks during development, we used rat cortical cultures in which *Ehmt1, Smarcb1, Mll3* or *Mbd5* were downregulated through RNA interference. Cultures were infected at day in vitro (DIV) 2 with lentiviruses expressing previously validated short hairpin RNAs (shRNAs) targeting *Ehmt1(25)* or newly designed shRNAs targeting *Smarcb1, Mll3* or *Mbd5.* Two independent shRNAs per gene were selected that reduced the respective expression levels by at least 50% (see(25) and Supplementary Figure S1a).

We recorded spontaneous electrical activity in all cultures by growing them on micro-electrode arrays (MEAs) (Supplementary Figure S1b). At the single channel level (black box, Supplementary Figure S1b), control neuronal networks (non-infected or GFP-infected) exhibit random events in the form of action potential spikes (highlighted in blue, Supplementary Figure S1b) and bursts (highlighted in pink, Supplementary Figure S1b). Together, these parameters are indicative of the overall spontaneous firing activity (i.e. firing rate, highlighted in grey, Supplementary Figure S1b). When bursts appear simultaneously in most of the channels (defined as 80% of the active channels), they form a synchronous event, called a network burst (green box, Supplementary Figure S1b). Typically, the pattern of activity in control neuronal networks develop following a stereotyped pattern (Supplementary Figure S1c) (26, 33). We found that early in development (i.e. DIV 10) neuronal networks displayed spontaneous electrophysiological activity comprised of random spikes and bursts. During the second week *in vitro* network bursts appeared, indicating that neurons start to functionally organize into a network. During development, the overall firing and network burst activity increased together with a reduction of the random spiking activity (Supplementary Figure S1d-f). Furthermore, the neuronal network synchronous activity developed from a stochastic towards a typical regular pattern. During the third week *in vitro*, the firing and network burst frequency plateaued and from this point on, the neuronal network activity remained stable. This stable state of activity indicates a functionally “mature” neuronal network.

To investigate whether KSS genes share a similar function during neuronal network formation and hence show common alterations at the neuronal network level when knocked down, we examined the electrophysiological activity of KSS gene-deficient networks and compared them to control cultures at DIV 10 (“immature state”) and DIV 20 (“ mature state”) (see raster plots in Figure 1a1-d1). We show that EHMT1-, SMARCB1 and MLL3-deficient neuronal networks were phenotypically similar during development. At DIV 10, these networks exhibited a higher level of random spiking activity, whereas the spike and network burst rates were similar compared to controls (Figure 1a_2_-a_4_, b_2_-b_4_, c_2_-c_4_). As the networks matured, the activity of EHMT1-, SMARCB1 and MLL3-deficient networks strongly increased. At DIV 20 these networks exhibited a higher level of activity (i.e. firing rate and/or network burst rate) compared to controls, indicating that the mature networks were in a hyperactive state (Figure 1a_5_, a_7_, b_5_, b_7_, c_5_, c_7_, Supplementary Table 1).

**Figure 1.**
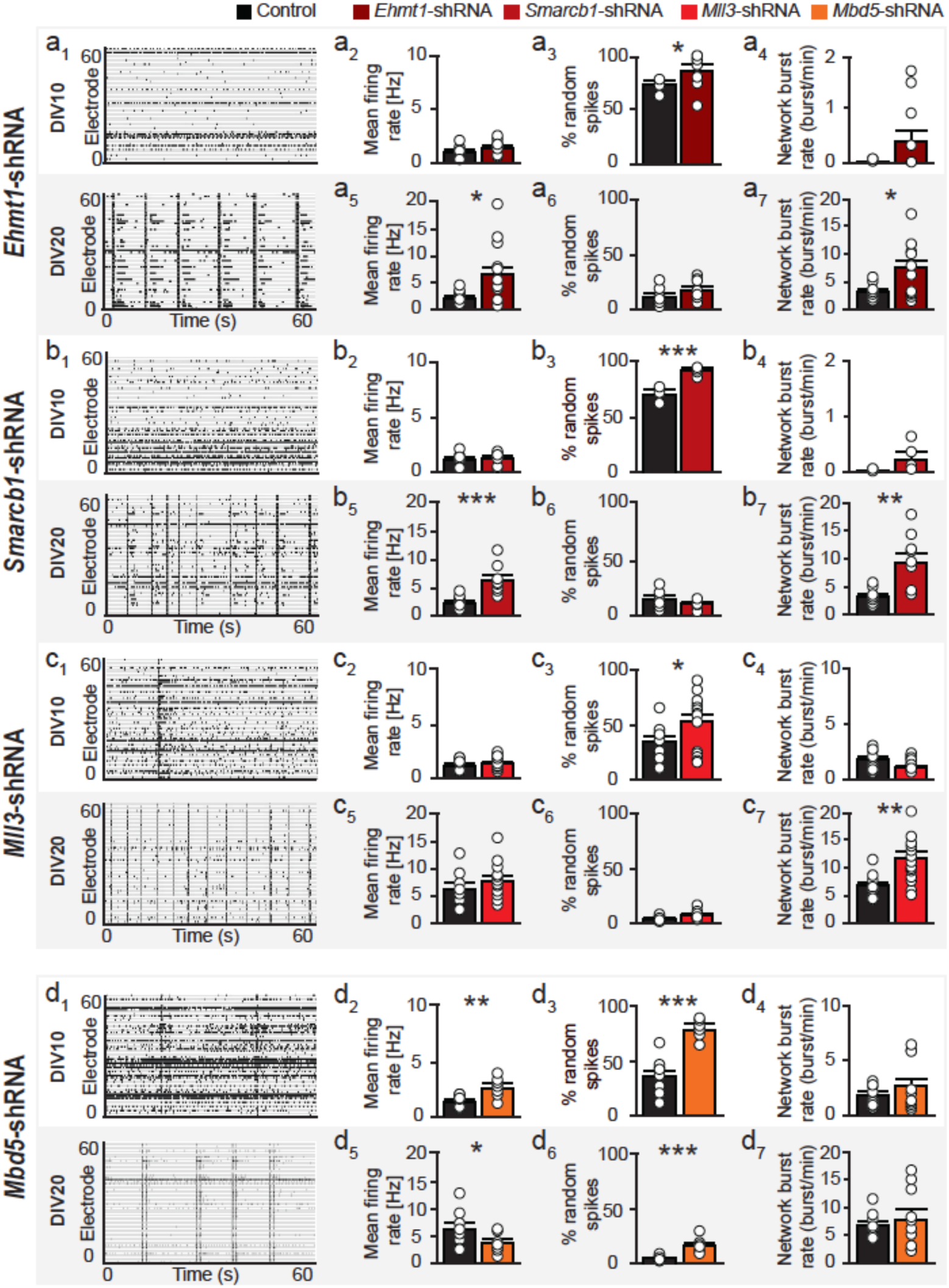
Network activity is altered in KSS-gene deficient cultures during development. a1-d1. Representative raster plots showing 60 s of electrophysiological activity recorded from KSS-deficient cultures at DIV10 and DIV 20. **a_2_-d_2_** MFR, **a_3_-d_3_** PRS and **a_4_-d_4_** NBR at DIV 10 in KSS-deficient cultures compared to control cultures. **a_5_-d_5_** MFR, **a_6_-d_6_** PRS and **a_7_-d_7_** NBR at DIV 20 in KSS-deficient cultures compared to control cultures (EHMT1-shRNA n=12; control: n=8; SMARCB1-shRNA: n=10; control: n=8; MLL3-shRNA: n=18; control: n=9; MBD5-shRNA: n=11; control: n=9). Sample size (n) indicates number of recorded MEA experiments over development. Data represent means ± SEM * p < 0.05, ** p < 0.01, *** p < 0.001 (parametric two-tailed T-test was performed between controls and KSS gene-deficient cultures). MFR: mean firing rate, PRS: percentage of random spike, NBR: network burst rate.

Although MBD5-deficient networks were also hyperactive, their developmental trajectory differed from the other KSS genes. At DIV 10, MBD5-deficient networks already showed an increase in overall activity expressed as mean firing rate (MFR, Figure 1d_2_), albeit with immature characteristics (i.e. more random spikes, Figure 1d_3_). The level of synchronous activity exhibited by controls and MBD5-deficient networks were similar at DIV 10 (Figure 1d_4_). Interestingly, whereas control neuronal networks increased their firing rate during development, MBD5-deficient neuronal networks did not. In fact, at DIV 20, MBD5-deficient networks exhibited less overall activity compared to controls (MFR, Figure 1d_5_) but with no differences in the network burst rate and a significantly higher number of random spikes (Figure 1d_6_, d_7_). This indicates that MBD5-deficient networks, although more active early in development, failed to organize properly by DIV 20.

Overall, we found prominent differences in the activity patterns exhibited by KSS gene-deficient neuronal networks compared to controls. Furthermore, our data indicate that shRNA-mediated knockdown of the KSS genes results in hyperactivity during development. This is particularly evident for EHMT1-, SMARCB1- and MLL3-deficient networks (Supplementary Figure S2c) and may suggest that a common pathophysiological mechanism leads to a global increase in neuronal excitability. The network phenotypes were all recapitulated with a second independent shRNA targeting each gene, indicating specificity (Supplementary Figure S1g, h).

### KSS gene deficiency alters neuronal network burst activity

Since our results showed that KSS gene-deficiency leads to network hyperactivity we next investigated if the typical pattern of network burst activity was also affected by studying network burst duration (NBD), network inter burst interval (NIBI) and network burst regularity. Whereas most of the network bursts (90.0%) in control neuronal networks lasted 200-600 ms (Figure 2a, f), we found that in KSS-deficient neuronal networks NBDs were differently distributed (Figure 2a-j). In particular, EHMT1-deficient networks showed NBDs longer than controls (48.2% of NBDs >600 ms, Figure 2g). SMARCB1-deficient networks exhibited NBDs both longer and shorter than controls (15.5% of NBDs >600 ms and 31.9% of NBDs <200 ms), indicated by multiple peaks in the distribution (Figure 2h). The NBDs distribution of MLL3- and MBD5-deficient networks was shifted to shorter durations compared to controls, indicated by the percentages of NBDs shorter than 600 ms (91.3%, 99.7% and 91.9% for control, MLL3- and MBD5-deficient networks, respectively; see distribution plot in Figure 2f, i, j).

**Figure 2.**
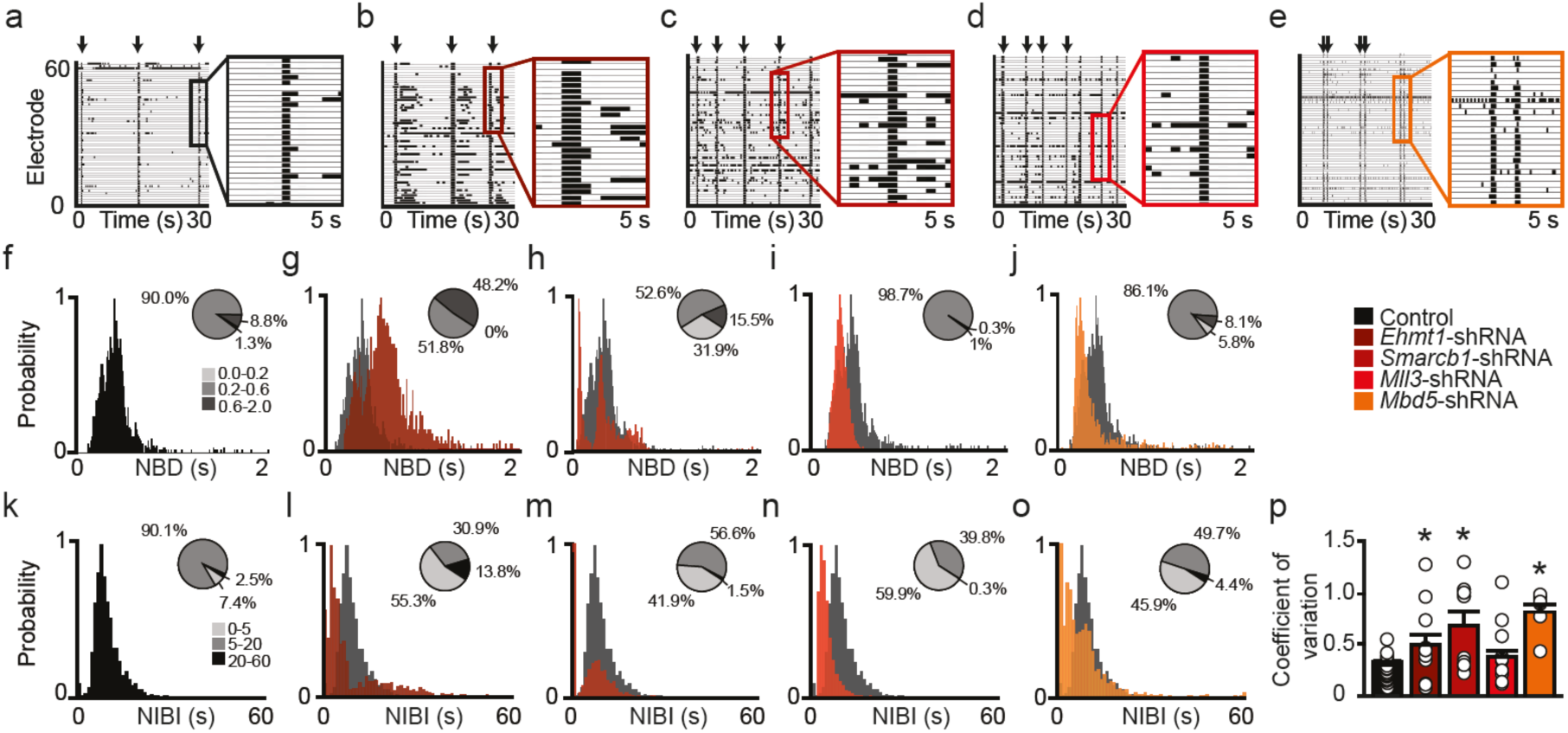
KSS gene deficiency alters neuronal network burst activity. **a-e.** Representative raster plots showing 30 s or recording of the electrophysiological activity of controls (**a**), EHMT1 (**b**) SMARCB1 (**c**), MLL3 (**d**) and MBD5 (**e**)-deficient cultures at DIV 20. Inset represents 5 seconds of recording displaying a network burst. **f-j.** Distribution of the duration of the network burst (i.e. NBD) exhibited by controls (**f**), EHMT1 (**g**) SMARCB1 (**h**), MLL3 (**i**) and MBD5 (**j**)-deficient networks (bin size of 10 ms). Pie diagrams display the percentage of network bursts with durations in three ranges: 0-0.2 s (light grey), 0.2-0.6 s (dark grey) and 0.6-2 s (black). **k-o.** Distribution of intervals between consecutive network bursts (i.e. NIBI) exhibited by controls (**k**), EHMT1 (**l**) SMARCB1 (**m**), MLL3 (**n**) and MBD5 (**o**) deficient networks (bin size of 1 s). Pie diagrams display the percentage of NIBI’s belonging to three intervals: 0-5 s (light grey), 5-20 s (dark grey) and 20-60 s (black). **p.** Coefficient of variation of the NIBI indicates the regularity of the network burst appearance in either EHMT1, SMARCB1, MLL3 and MBD5-deficient cultures as compared to controls at DIV 20. Sample seize n=17 for controls, n=12 for *Ehmt1-shRNA*, n=10 for *Smarcb1-shRNA*, n=18 for *Mll3*shRNA and n=11 for *Mbd5-shRNA* for all displayed measurement. Data represent means ± SEM. * p < 0.05 (parametric two-tailed T-test was performed between controls and KSS gene-deficient cultures).

Then, we studied how each KSS gene knockdown affected the NIBI. The majority (90.1%) of NIBIs in control neuronal networks occurred within a range of 5-20 s (Figure 2k). In contrast, we observed that KSS genes-deficient networks showed NIBIs shorter than controls (55.3%, 41.9%, 59.9% and 45.9% of NIBIs <5 s for EHMT1-, SMARCB1-, MLL3- and MBD5-deficient networks respectively; Figure 2l-o).

Finally, we investigated whether KSS genes knockdown affected the typical regular network burst pattern exhibited by control neuronal networks. To determine the regularity, we computed the coefficient of variation of the NIBIs. We found that all KSS-genes deficient networks, except MLL3, exhibited an irregular network burst pattern, as indicated by the higher coefficient of variation of the NIBIs compared to controls (Figure 2p).

In summary, our data indicate that KSS gene-deficient neuronal networks become hyperactive during development and showed impairments in the pattern of network burst activity late in development.

### EHMT2-deficient networks show a different phenotype compared to KSS gene-deficient networks

EHMT2 is a paralogue of EHMT1, but has not been associated with KSS or any other NDD. To investigate whether the network phenotypes are specific to KSS gene deficiency, we knocked down *Ehmt2* using validated shRNAs(25) in developing neuronal cultures. In contrast to KSS gene-deficient networks, EHMT2-deficient neuronal networks exhibited significantly lower mean firing rates both at DIV10 and DIV20 (Supplementary Figure S2). This further confirms our previous observations that loss of EHMT1 or EHMT2 in neurons can generate distinct phenotypes(23, 25).

### Deficiency of KSS genes leads to increased neuronal excitability

The increased neuronal network activity we found at DIV 20 might be caused by altered intrinsic neuronal parameters resulting in hyperexcitability of the individual neurons and/or changes in extrinsic parameters related to synaptic signaling. Supporting this notion, intrinsic parameters linked to neuronal excitability have recently been shown to be regulated, at least in part, by epigenetic modifications via DNA methylation(34). Using whole-cell patch-clamp recordings of individual neurons at DIV 20, we measured intrinsic passive and active electrophysiological properties (Figure 3a-f, Supplementary Table 2).

**Figure 3.**
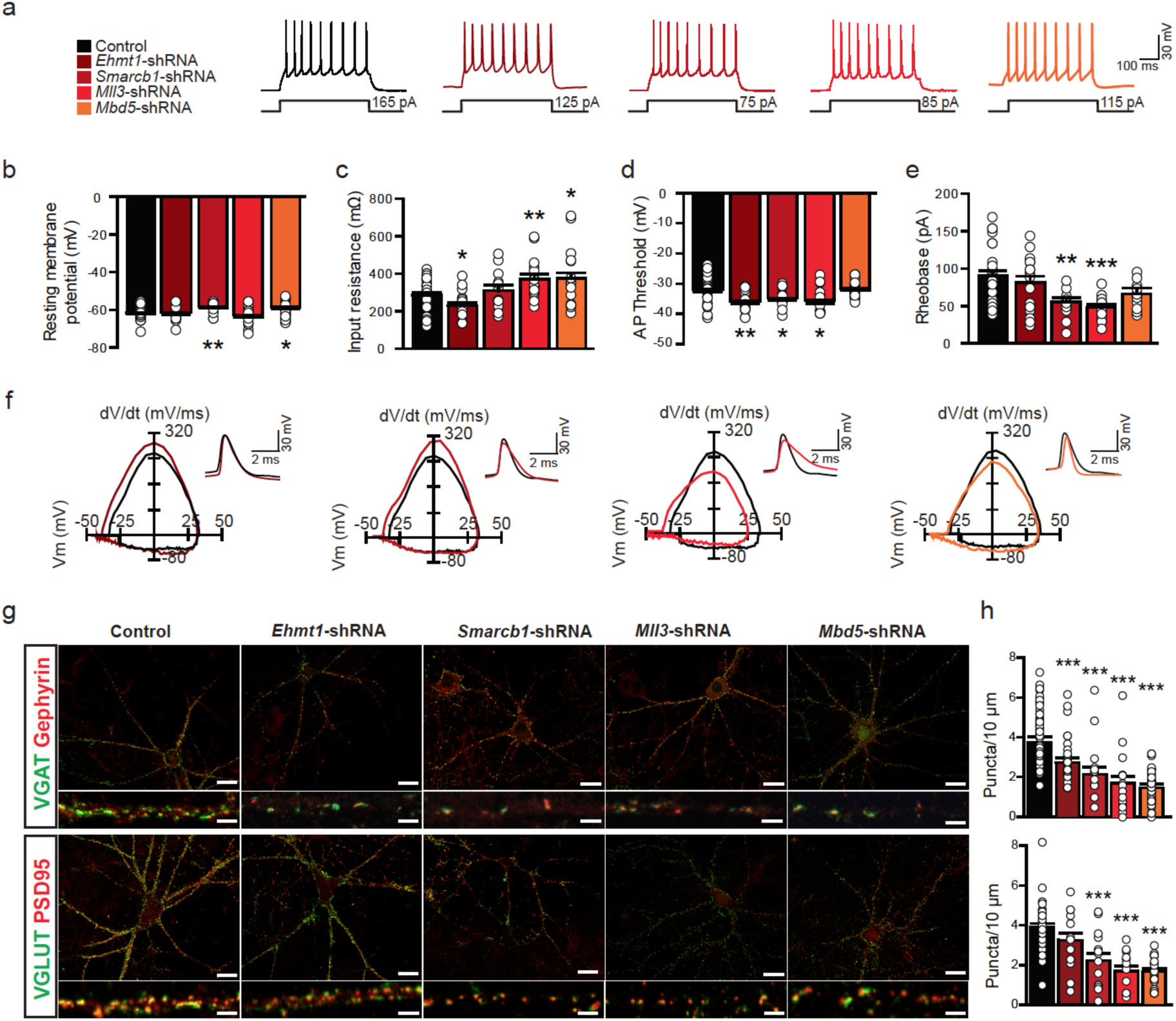
Increased excitability and altered excitatory/inhibitory synaptic inputs in neurons of KSS gene-deficient networks. **a.** Representative firing patterns of neurons in KSS gene-deficient cultures. **b-e.** Passive intrinsic properties (**b-c**) and active intrinsic properties (**d-e**) of neurons in KSS gene-deficient networks at DIV20. (n=28 for controls, n=18 for *Ehmt1-shRNA*, n=13 for *Smarcb1-shRNA*, n=16 for *Mll3* shRNA and n=19 for *Mbd5-shRNA* for all displayed measurements). Sample size (n) indicates number of cells per genotype. **f.** Representative outline of a single action potential waveform and phase-plot. **g.** Immunocytochemical analysis of inhibitory (VGAT and Gephyrin co-localized puncta) and excitatory (VGLUT and PSD95 co-localized puncta) synapses representing in control and KSS gene-deficient cultures at DIV 20. **h.** Quantification of number of inhibitory and excitatory synapses per 10 µM dendrite in all KSS gene-deficient cultures. Scale bar represents 10 µM (upper panel) and 2 µM (lower panel). Data represent means ± SEM. * p < 0.05; ** p < 0.01; *** p < 0.001 (parametric two-tailed T-test was performed between controls and KSS gene-deficient cultures).

In EHMT1-deficient neurons we found a hyperpolarizing shift of the action potential (AP) threshold combined with unaltered resting membrane potentials (V_rmp_) (Figure 3b, d). At standard holding potentials (−60 mV) however these changes did not result in a reduction in the AP firing rheobase (Figure 3a, e), since the EHMT1-deficient neurons also showed lower membrane input resistances (R_in_, Figure 3c).

Similar to EHMT1 knockdown, SMARCB1- and MLL3-deficient neurons both showed a hyperpolarizing shift of the AP threshold at unaltered (MLL3) or depolarized V_rmp_ (SMARCB1) (Figure 3b, d). In addition, SMARCB1-deficient neurons showed an unchanged R_in_ and MLL3-deficient neurons showed an increased R_in_ (Figure 3c). These alterations may underlie our finding that both of these KSS-gene deficiencies share a reduced AP firing rheobase (Figure 3a, e), which supports increased neuronal excitability.

Neurons in MBD5-deficient networks were the only ones that showed no change in the AP threshold (Figure 3d). Even though at −60 mV the rheobase remained unchanged (Figure 3e), the generally depolarized V_rmp_ (Figure 3b) in combination with an increased R_in_ (Figure 3c) still implies an increased excitability of MBD5-deficient neurons due to a higher responsiveness to incoming excitatory (depolarizing) synaptic current at V_rmp_.

For all tested KSS genes, we thus found changes in intrinsic properties that directly (AP threshold) or indirectly (V_rmp_, R_in,_, τ) affect the generation of APs. Therefore, we compared the AP waveforms across genotypes (Figure 3f, Supplementary Table 2). Whereas APs generated by EHMT1-deficient neurons showed no significant changes in their AP waveform, SMARCB1- and MLL3-deficient neurons both showed broader APs, mediated by a slower rising phase (i.e. rise-time, MLL3-deficient neurons only) and/or slower repolarisation phase (i.e. decay time, SMARCB1- and MLL3-deficient neurons). Contrasting these, APs in MBD5-deficient neurons were significantly shorter, due to a faster repolarisation phase.

Taken together these results indicate that alterations in intrinsic passive and active properties in KSS-gene deficient neuronal networks is genotype specific, but as a whole imply different levels of increased neuronal excitability.

### KSS-deficiency leads to altered excitatory and inhibitory synaptic inputs

In addition to increased intrinsic excitability, the hyperactivity observed in KSS gene-deficient networks could also be explained by a change in excitatory/inhibitory (E/I) balance. To investigate this, we measured synaptic properties in our cultures, with and without KSS gene knockdown. We first measured miniature inhibitory postsynaptic currents (mIPSCs) in EHMT1-deficient networks at DIV 20 (Supplementary Figure S3). We found a significant reduction in mIPSC frequency, but not in mIPSC amplitude, when compared to control cultures. We previously showed that knockdown of EHMT1 did not affect miniature excitatory postsynaptic current (mEPSC) frequency or amplitude in rat neuronal networks(25). Therefore, our combined results suggest that that the E/I balance is shifted in favor of excitation due to reduced inhibitory synaptic input.

Because mIPSC frequency is known to correlate with the number of synapses and release probability of a neuron, we counted the number of synapses in both control and KSS gene-deficient neuronal networks (Figure 3g, h). First, we quantified the number of inhibitory synapses on individual dendrites by counting the number of colocalizing presynaptic vesicular GABA transporter (VGAT) and postsynaptic Gephyrin puncta. We found a significant reduction in the number of inhibitory synapses for all KSS genes when compared to control cultures (Figure 3g, h). Thus, KSS gene deficiency likely has a strong effect on the formation and/or maintenance of inhibitory synapses. Next we quantified excitatory synapses, by counting the number of presynaptic vesicular glutamate transporter (VGLUT) and the postsynaptic density-95 protein (PSD95) puncta colocalizing. We found a significant reduction in the number of excitatory synapses in SMARCB1-, MLL3- and MBD5-deficient neuronal networks, but we found no changes in the EHMT1-deficient networks (Figure 3g, h), in line with our previous reports(25, 26).

In summary, we show that in EHMT1-deficient networks the E/I balance is strongly shifted to increased excitation due to reduced inhibitory synaptic input. SMARCB1-, MLL3- and MBD5-deficient cultures, showed a reduction in inhibitory input, which was also accompanied with a reduction in excitatory input.

### KSS deficiency causes deregulation of genes controlling neuronal and synaptic processes

Next, we investigated the molecular changes that could underlie the hyperactivity that we observed in KSS gene-deficient networks. To address this, we performed RNA sequencing on DIV 20 neuronal networks after KSS gene knockdown.

Using DESeq2(35), in all KSS gene-deficient cultures, we detected differentially expressed (DE) genes (q value < 0.1), as compared to control cultures. Knockdown of *Ehmt1* gave rise to more up-regulated than downregulated genes. In contrast, we detected more downregulated genes than up-regulated genes in SMARCB1-, MLL3- and MBD5-deficient cultures (Figure 4a and Supplementary Table 3). This observation indicates that SMARCB1, MLL3 and MBD5 regulate gene transcription in an opposite direction, as compared to EHMT1. This is consistent with that SMARCB1 and MLL3 are known to be transcriptional activators and EHMT1 to be a transcriptional repressor(36, 37). In addition, EHMT1- and MBD5-deficient cultures showed a lower number of total DE genes compared to SMARCB1- and MLL3-deficient cultures (Figure 4a). To gain an overview of the global gene expression pattern, we performed a principal component analysis (PCA) on DE genes obtained from all pair-wise comparisons (3083 genes, Figure 4b). PCA allowed discrimination of KSS gene-deficient samples, with DE genes of SMARCB1- and MLL3-deficient cultures being close to each other, on the opposite end of DE genes of EHMT1-deficient cultures. The DE genes of MBD5-deficient cultures were found to be closest to the control. Furthermore, the comparisons of DE genes between each pair of the KSS knockdown using the scatter plot analysis (Supplementary Figure S4) showed that DE genes of SMARCB1- and MLL3-deficient cultures had the highest correlation (r^2^ = 0.90). These data indicate that knockdown of different KSS genes resulted in distinguishable gene expression patterns, where those of *Smarcb1* and *Mll3* were most similar and that of *Ehmt1* was most different. Interestingly, GO annotation of DE genes detected from EHMT1-, SMARCB1- and MLL3-deficient cultures shared high similarity in their associated biological functions (biological processes, BP), in particular, in ion transmembrane transport and chemical synaptic transmission (Figure 4c). DE genes detected in MBD5-deficient cultures were associated with apparently different biological functions including cell adhesion, nucleosome assembly and protein translation. However, GO annotation assessed for cellular components (CC) revealed that DE genes detected from all knockdown conditions were very similar, mainly associated with axon, dendrite, synapse and postsynaptic density (Figure 4d). These results indicate that the similar neuronal structures are affected by KSS gene knockdown through distinct molecular mechanisms. A closer examination of DE genes that are known to play roles in synaptic and ion channels functions showed that most of these genes were affected by knockdown of all four KSS genes, but the regulation was different, with opposite expression patterns of *Smarcb1/Mll3* and *Ehmt1* knockdown, and a unique pattern of *Mbd5* knockdown (Figure 4e). In addition, we identified 34 DE genes represented in all knockdown conditions (Supplementary Figure S5a). Also here the expression patterns of these 34 DE genes were mostly in opposing directions between SMARCB1/MLL3- and EHMT1-deficient cultures (Supplementary Figure S5b, Supplementary Table 4). Remarkably, this small number of 34 DE genes revealed an enrichment of GO terms related to learning, memory, neurons and dendrites (Supplementary Figure S5c, d). Of interest, many of the 34 DE genes have previously been associated with cognitive disorders, seizures or epilepsy, ASD, motor abnormalities, and sleep disturbances (Supplementary Table 5), which is a constellation of symptoms seen in KSS.

Taken together, these data show that KSS target genes share similar functions in regulating neuronal structures and activity, with a prominent enrichment for genes that directly affect neuronal excitability (e.g. potassium and sodium channels) and synaptic function, including several GABA and glutamate receptors (Figure 4e). However KSS genes regulate distinct sets of individual target genes through different transcriptional or functional mechanisms. The difference at the functional level was most apparent for MBD5, which was consistently the most dissimilar of the KSS genes, from the functional level (Figure 1, Supplementary Figure S2c) to gene expression level.

**Figure 4.**
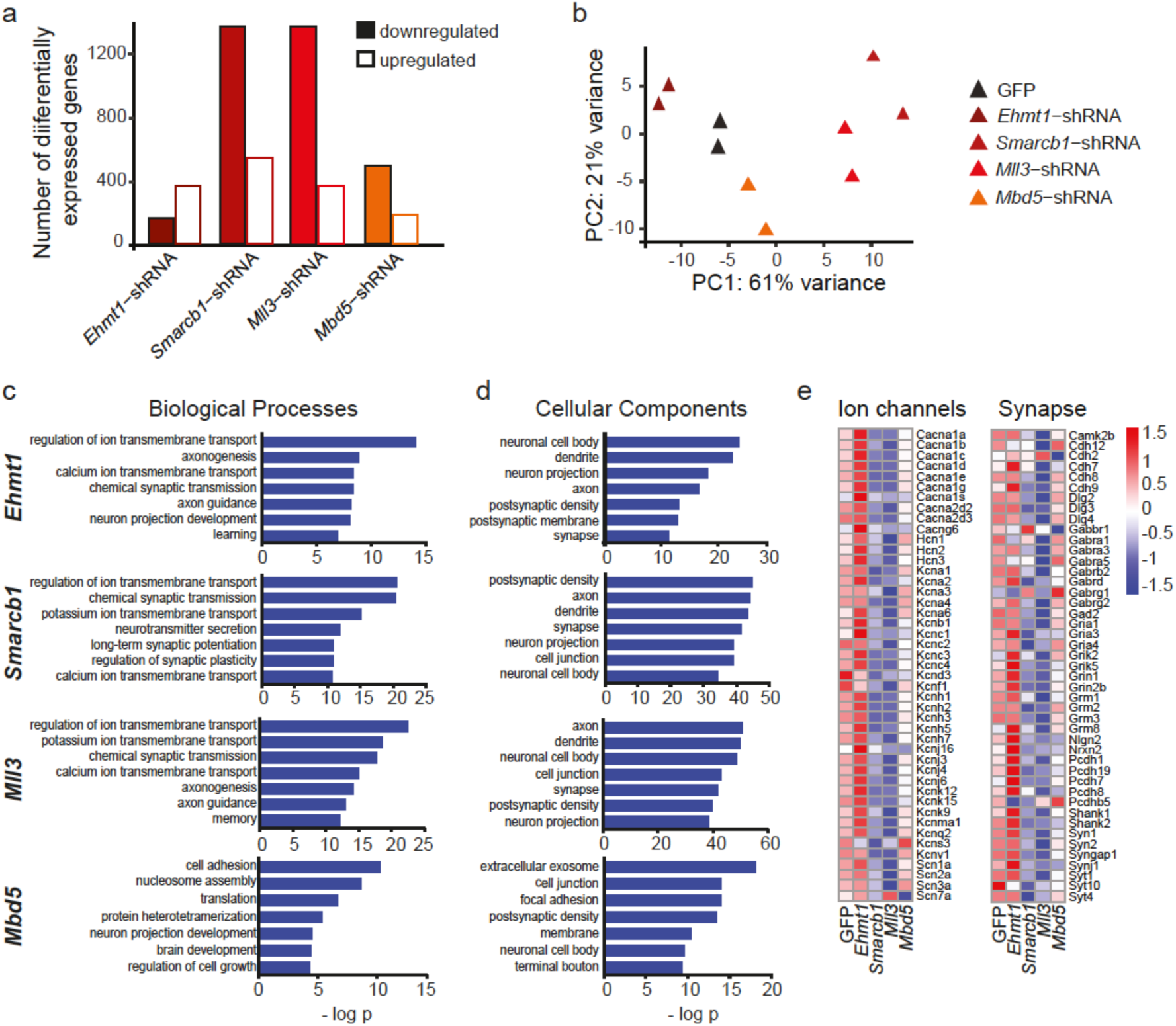
KSS deficiency caused deregulation of genes involved in neuronal and synaptic function. **a.** Number of differentially expressed genes (DE) in the four knockdown conditions, as compared to GFP knockdown. **b.** PCA plot using DE genes (q < 0.1) of four knockdown samples and GFP knockdown control sample (in duplicates). **c.** Top seven Gene Ontology (GO) annotation terms in the category of ‘biological processes’ detected from the DE genes (q < 0.1). **d.** Top seven GO terms in the category of ‘cellular components’ detected from the DE genes (q < 0.1). **e.** Heatmap of relative expression (Z scores) of a selection of DE genes (q < 0.1) that are known to play roles in ion channels and synapse detected in the four knockdown samples, as compared to the control (GFP knockdown). The scale of the Z scores is indicated from −1.5 (blue) to 1.5 (red).

### Increased cell excitability and reduced inhibition in *Ehmt1^+/-^* mice

Having established that loss of EHMT1 leads to increased cell excitability and reduced synaptic inhibition *in vitro*, we aimed to corroborate these results by measuring intrinsic and synaptic properties in acute hippocampal brain slice preparations of *Ehmt1^+/-^* mice.

First, we examined the development of synaptic inputs by recording mIPSCs and mEPSCs at postnatal day (P) 7, P14 and P21 in *Ehmt1^+/+^* and *Ehmt1^+/-^* mice, revealing a reduction in mIPSC amplitudes at all investigated time points (Figure 5a, b and Supplementary Figure S6b, e) in CA1 pyramidal neurons. We found an increase of mIPSC frequency at P7, but a strong reduction at P21 (Figure 5c and Supplementary Figure S6c, f) leading to a general reduction of inhibitory connectivity at P21, consistent with our observation in dissociated rat cortical neurons. Recording of the paired pulse ratio (PPR) at P21 revealed an increased PPR specifically at 50 ms inter-stimulus interval (ISI) following stimulation in *stratum radiatum* (Figure 5d but not *stratum oriens* (Supplementary Figure S6m, n), indicating that these interneurons have a reduced probability of release onto CA1 pyramidal cells. Interestingly, mEPSC amplitude and frequency were unaltered between *Ehmt1^+/+^* and *Ehmt1^+/-^* mice (Figure 5e-g and see Supplementary Figure S6g-l). In addition, recording of the PPR following stimulation of the Schaffer collaterals, the main excitatory input to CA1 pyramidal neurons, showed no changes in the probability of release at P21 (Figure 5h). These data confirm our in vitro data and suggest that EHMT1 plays a role in controlling E/I balance by regulating inhibitory inputs onto CA1 pyramidal cells. This is in line with the expression pattern of EHMT1, which next to excitatory neurons(21), we also found to be expressed in both parvalbumin and somatostatin positive cells (Supplementary Figure S6o).

**Figure 5.**
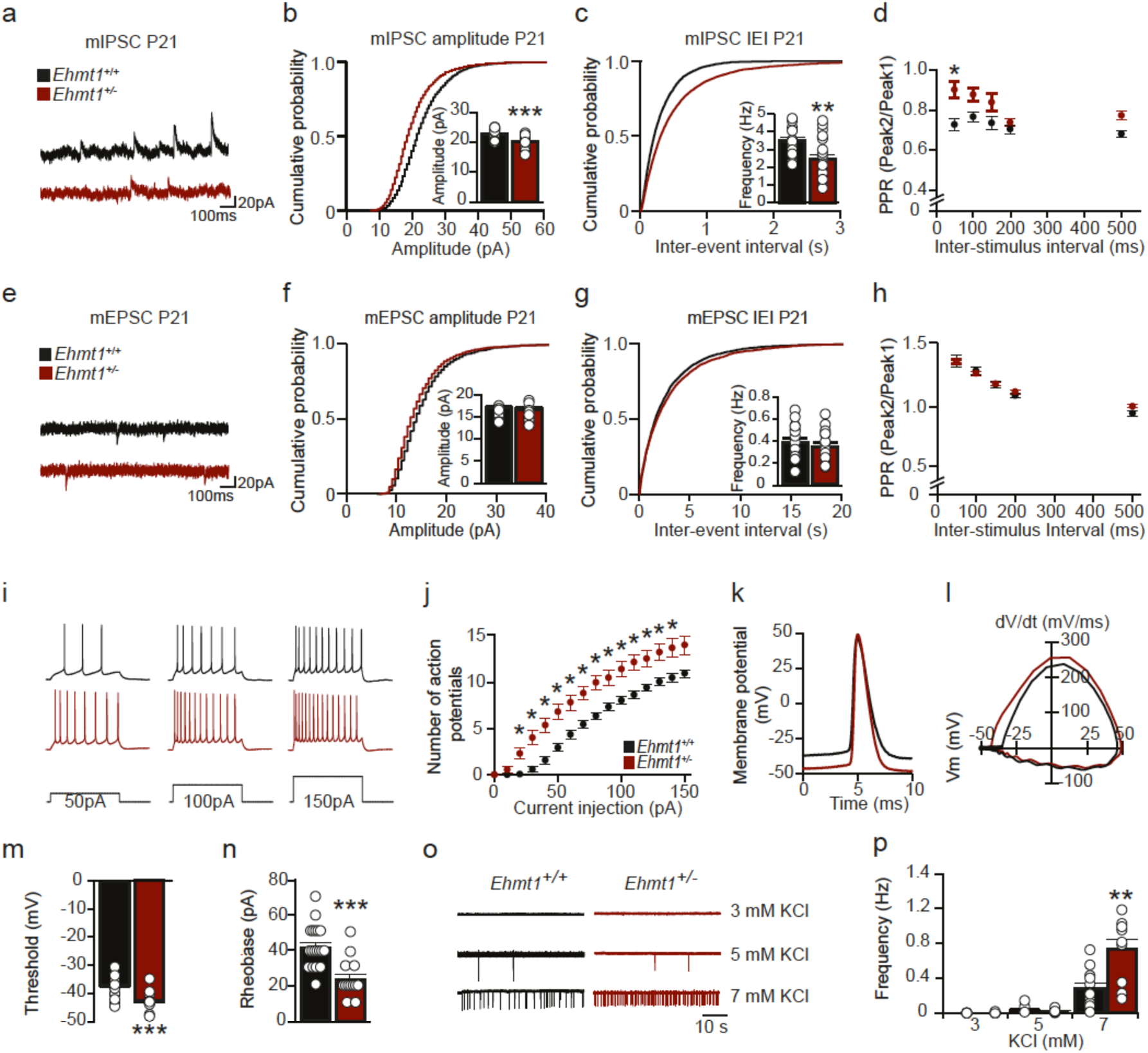
Increased synaptic inhibition and intrinsic properties in *Ehmt1* mice. **a.** Example traces of mIPSC recordings from *Ehmt1^+/+^* and *Ehmt1^+/-^* mice at P21. **b-c**. Quantification of mIPSC amplitude (**b**) and frequency (**c**) from *Ehmt1^+/+^* and *Ehmt1^+/-^* mice at P21 (n=19 for *Ehmt1^+/+^* and n=22 for *Ehmt1^+/+^* for both parameters) **d.** PPR responses following stimulation in SR at P21 (n=8 for *Ehmt1^+/+^* and n=8 for *Ehmt1^+/-^*). **e.** Example traces of mEPSC recordings from *Ehmt1^+/+^* and *Ehmt1^+/-^* mice at P21. **f-g.** Quantification of mEPSC amplitude (**f**) and frequency (**g**) from *Ehmt1^+/+^* and *Ehmt1^+/-^* mice at P21 (n=17 for *Ehmt1^+/+^* and n=15 for *Ehmt1^+/-^* for both parameters). **h.** PPR responses following stimulation of the Schaffer collaterals at P21 (n=13 for *Ehmt1^+/+^* and n=17 for *Ehmt1^+/-^*). **i.** Firing patterns of *Ehmt1^+/+^* and *Ehmt1^+/-^* mice pyramidal neurons in response to current injections of 50 pA, 100 pA and 150 pA at P21. **j.** Quantification of intrinsic excitability of *Ehmt1^+/+^* and *Ehmt1^+/-^* neurons at P21 (n=19 for *Ehmt1^+/+^* and n=13 for *Ehmt1^+/-^*). **k.** Example outlines of the action potential waveforms of *Ehmt1^+/+^* and *Ehmt1^+/-^* neurons. **l.** Phase-plot of the first AP at rheobase of *Ehmt1^+/+^* and *Ehmt1^+/-^* neurons. **m-n.** Quantification of the threshold (n=19 for *Ehmt1^+/+^* and n=13 for *Ehmt1^+/-^*) and rheobase (n=22 for *Ehmt1^+/+^* and n=14 for *Ehmt1^+/-^*) of *Ehmt1^+/+^* and *Ehmt1^+/-^* pyramidal neurons. **o.** Example traces of spontaneous AP firing in different concentration of KCl. **p.** Quantification of spontaneous AP firing upon KCl treatment (n=5 (3 mM), n=6 (5 mM) and n=11 (7 mM) for *Ehmt1* and n=6 (3 mM), n=7 (5 mM) and n=11 (7 mM) *for Ehmt1* mice). * p < 0.05; ** p < 0.01; *** p < 0.001. IEI: Inter-event interval. PPR: Paired-pulse ratio. Data represents mean ± SEM, Two-tailed Student’s t-test, PPR was Bonferroni corrected.

In analogy to the primary neuronal cultures, we then investigated the intrinsic excitability of CA1 pyramidal neurons by means of their intrinsic electrophysiological properties. An input/output curve with increasing amounts of injected current revealed an increased excitability of CA1 pyramidal neurons (Figure 5i, j), which was accompanied by a reduced rheobase (Figure 5n). This reduction of the rheobase can be specifically attributed to a hyperpolarization of the AP threshold (Figure 5k-m), since other intrinsic properties remained unchanged (Supplementary Table 6). These results indicated that CA1 pyramidal cells were intrinsically more excitable in *Ehmt1^+/-^* mice compared to *Ehmt1^+/+^* mice.

Since CA1 pyramidal cells show a reduced inhibitory synaptic connectivity and an increased intrinsic excitability, we hypothesized that these cells should display higher level of spontaneous spiking activity. To investigate this, we performed cell-attached patch clamp recordings of CA1 pyramidal neurons to record basal AP frequency. In standard recording solution (3 mM KCl), CA1 pyramidal cells are inactive, as reported previously^12^. Elevating KCl concentration to 7 mM resulted in AP firing in all recorded cells, and revealed a higher AP frequency in *Ehmt1^+/-^* compared to *Ehmt1^+/+^* mice (Figure 5o, p). These data indicate that the combination of reduced inhibitory synaptic inputs and an increased intrinsic excitability may result in increased basal activity of CA1 pyramidal neurons in *Ehmt1^+/-^* mice.

## Discussion

In this study, we took advantage of the genetic heterogeneity observed in KSS (caused by *EHMT1, SMARCB1, MLL3 or MBD5* loss of function) to investigate whether the effects of those mutations share similar patterns of molecular, cellular and/or neuronal network (dys)function. We used a combination of gene expression analysis, single-cell patch clamp and neuronal network recordings to investigate and compare the consequences of KSS gene deficiency. We demonstrated that the effects from partial loss of each KSS gene overlapped at the network level and were characterized by increased cellular excitability and network hyperactivity. At the molecular level, gene networks that were altered converged on biological pathways related to potassium channels and synaptic communication.

### KSS gene-deficient networks share a common mode of failure

Previously we showed that EHMT1 deficiency transiently delays the appearance of spontaneous network activity, eventually resulting in an irregular network bursting pattern(26). Here, we show that following excessive random spiking activity in immature cultures, the irregular network burst pattern is generally accompanied by a more frequent network burst rate (i.e. network hyperactivity) in mature EHMT1-deficient networks. Interestingly, the network phenotypes after loss of the other KSS genes showed striking similarities. Knocking down SMARCB1, MLL3 or MBD5 also resulted in an irregular network burst pattern and/or hyperactivity. Our results therefore imply that the KSS gene-deficient networks share a common mode of failure when establishing network communication (Supplementary Figure S2c). Of note, this implied common mode of failure was reflected in different compositions of altered parameters of network activity in a genotype specific manner. For example, hyperactivity was observed as an increase in both firing and network burst rate in EHMT1- and SMARCB1-deficient neuronal networks, while only one of the two parameters was altered in MLL3-(i.e. network burst rate) and MBD5-(i.e. firing rate) deficient neuronal networks. The developmental trajectory for MBD5-deficient networks was representative for the genotype specific differences too.

While the other knockdowns showed hyperactivity late in development, MBD5-deficient networks were excessively active at an early, immature stage (DIV10). The functional convergence we observed at the neuronal network level is in line with a recent study in *Drosophila* showing similar deficits in short-term memory between flies lacking EHMT1 and MLL3 in mushroom bodies(27). In general, hyperactivity in networks can be mediated by two major factors: a) changes in synaptic signaling between neurons resulting in altered E/I-balance; and b) changes in intrinsic electrophysiological properties of the neurons within the networks resulting e.g. in hyperexcitability(38). At the single cell level, we found in SMARCB1-, MLL3- and MBD5-deficient cultures a strong reduction in both excitatory and inhibitory synaptic inputs. The reduction (~50%) was similar for excitatory and inhibitory synapses, suggesting that the E/I balance was not changed by those knockdowns. EHMT1-deficient cultures, however, showed a strong decrement in inhibitory synaptic input, without excitatory synaptic input being affected. This was the case both *in vitro* as well as in the *Ehmt1^+/-^* mice. Indeed, we found mIPSC amplitude to be decreased at all investigated time points in the *Ehmt1^+/-^* mice, whereas the mIPSC frequency was strongly reduced at P21. This effect on frequency can be explained by fewer inhibitory synapses and by a reduced release probability of inhibitory synapses following stimulation in *stratum radiatum*, but not *stratum oriens.* The specific effect of loss of *Ehmt1* on inhibition is relevant because imbalanced E/I is associated with ASD in humans and rodent models(39–42). In particular, a loss in the efficiency of inhibitory synaptic strength has been observed in many NDDs, including Rett syndrome and Fragile X syndrome(43–47). The changes in excitatory and inhibitory inputs observed in KSS gene-deficient neurons imply alterations in proteins directly or indirectly linked to synapse function. In line with this concept, for all KSS gene knockdowns we found a multitude of DE genes linked to both glutamatergic and GABAergic synaptic transmission, including up- and downregulation of glutamate and GABA receptors, adhesion molecules and postsynaptic density proteins. Presumably, multiple genes are responsible for the observed functional changes and, at the same time, a portion of the underlying transcriptional changes could be subtractive or antagonistic in nature, resulting in limited functional consequences for synaptic transmission. Further studies would be required to identify direct versus indirect targets, for example through the identification of the underlying epigenetic changes. In addition to the shift in E/I balance that we found in EHMT1-deficient neurons, neuronal hyperexcitability could contribute to increased and/or irregular network burst activity in KSS gene-deficient networks(38). Enhanced neuronal excitability can be mediated by single or combinations of passive as well as active intrinsic properties. These properties encompass increased membrane input resistance or a hyperpolarized shift in the threshold for generating action potentials, particularly in combination with a depolarized membrane potential. Indeed, at the single cell level across all four investigated KSS genes we found changes in the intrinsic neuronal properties that imply increased excitability. However, the increased excitability in KSS-deficient neurons was genotype specific, both in terms of contributing properties and extent of increase (i.e. mild for EHMT1- and MBD5-deficient networks and more pronounced for SMARCB1- and MLL3-deficient ones). Furthermore, we found robust genotype specific alterations in the action potential kinetics, ranging from relatively slow (SMARCB1 and MLL3) to fast (MBD5) repolarization kinetics. The various changes in passive and active parameters that result in increased neuronal excitability in KSS gene-deficient networks suggest that ion channel expression is altered. In particular, reduced expression of different classes of voltage gated potassium (K_V_) channels, such as K_v_1.1 (KCNA1) or K_v_2.1 (KCNB1), and/or increased expression of voltage gated sodium (Na_V_) channels, such as SCN1A, SCN2A or SCN3A, has been shown to increase somatodendritic excitability and action potential kinetics(48)(49)(50). Furthermore, dysregulation or dysfunction in several classes of K_v_ and Na_v_ channels, including those mentioned above, have been found to be associated with NDDs(51), epilepsy and ASD(52). Strikingly, in all KSS gene-deficient networks we found altered regulation of a battery of genes coding for different types of ion channels, including several classes of K_v_ and Na_v_ channels. However, the diversity and similarities in the intrinsic electrophysiological parameters is also reflected by changes in gene expression. Neurons haploinsufficient for the repressive regulator EHMT1 almost exclusively show upregulation of genes coding for K_v_ and Na_v_ channels, whereas neurons deficient for SMARCB1 and MLL3 consistently show downregulation in these genes. The opposing up and down-regulation of genes for ion channels illustrates that the detected hyperexcitability is likely to be the combined consequence of complex changes in ion channel composition that can either be dominated more by increased Na_v_ channel expression (EHMT1), more by reduced K_v_ channel expression (SMARCB1 and MLL3) or by a complex combination of both (MBD5).

### Molecular convergence in KSS

An intriguing finding that we uncovered by comparing RNA-Seq of each knockdown is a set of 34 commonly dysregulated transcripts. The list comprises genes that are associated with cognitive disorders, epilepsy or ASD, and most code for proteins involved in synaptic function or ion channels (see Supplementary Table 5). Interestingly, four of these genes have been identified as hub genes in co-expression networks analyzed from the cortical tissue of ASD patients: *Scamp5, Slc12a5, SynJ1* and *Unc13a* (4)(53)(54). Another transcript, *Adcy1*, which is strongly upregulated by *Ehmt1* knockdown was recently found to be upregulated in *Fmr1 KO* mice, a model of Fragile X syndrome, and specifically reducing *Adcy1* expression ameliorated the autistic behaviors(55). *Apc* is a negative regulator of the canonical Wnt/β-catenin signaling pathway and its expression is also significantly altered by all KSS gene knockdowns. One study showed that conditional knockout of *Apc* in forebrain neurons causes autistic-like features in mice(56).

The majority of genes in the list are upregulated after *Ehmt1* knockdown but downregulated by *Smarcb1, Mll3* or *Mbd5* loss of function. Accordingly, EHMT1 enzymatic activity generally represses transcription while SMARCB1 and MLL3, and according to our data MBD5, function as activators. The divergent effects on mRNA expression may explain some phenotypic differences we observed. There are four transcripts upregulated by *Ehmt1* knockdown that are directly involved in high-frequency neuronal activity: the protein products of *Scamp5* and *Unc13a* (aka, Munc13-1) maintain high rates of vesicular endo- and exocytosis(57, 58), while *Scn8a* codes for the sodium channel Na_V_1.6, whose channel properties support high firing rates(59). Finally, *Grin1* is an interesting transcript since enhancing NMDAR activity has been directly implicated in lengthening burst duration(38). We speculate that manipulating only these genes in control neurons may recapitulate the burst pattern or conversely restore the phenotype observed in EHMT1-deficient networks.

The 34 overlapping genes may be inexorably linked (representing a neuronal co-expression module) and at least some may be direct, common targets of the pathogenic epigenetic modulators found in KSS. It would therefore be useful to decipher the epigenetic marks that control expression, especially since it is not clear which transcriptional changes are causal versus those that are collateral. Importantly, our data suggest that the molecular pathophysiological mechanism underlying KSS may not depend on whether the parallel transcriptional changes are a gain or loss. Instead, the implication is that a dysfunction in dynamic transcriptional regulation during development leads to disease and consequently hinders proper neuronal specialization or cortical patterning, as has been suggested to occur in autism(4).

In this study we show that KSS genes mainly converge at the neuronal network level. Underlying this functional convergence, we found that common biological pathways affected which related to common ion-channel expression and synaptic communication, albeit we clearly observed distinct patterns of deregulation for each of the KSS genes. As neuronal network behavior and organization are defined by its molecular and functional history, our measurements in neuronal cultures provide a powerful tool to investigate not only the pathophysiological mechanisms underlying KSS, but also those of NDDs and other neurological disorders.

## Material and methods

### RNA interference

For RNAi knockdown experiments, DNA fragments encoding short hairpin RNAs (shRNAs) directed against rat *Ehmt1*(25), *Ehmt2*(25), *SmarcB1, Mll3* or *Mbd5* mRNA were cloned into the pTRIPΔU3-EF1α-EGFP lentiviral vector(60, 61). Used hairpin sequences encompassed: *Smarcb1* hp#1: GGAGATTGCCATCCGAAAT, *Smarcb1* hp#2: GCCCTCCTTCAGCACACAT, *Mll3* hp#1:

GGCCTCCATTCACACCAAT, *Mll3* hp#2: GGCCAAGACCCTGCTGTAA, *Mbd5* hp#1:

CCGGAAATGGTTCTGTAAAGAGT, *Mbd5* hp#2: CTGAAGGACACAGCACTTTAAAC. Empty vector expressing GFP only was used as control vector. Lentiviral particles were prepared, concentrated, and tittered as described previously(60). In brief, lentiviruses were generated by co-transfecting the transfer vector, the psPAX2 packaging vector (Addgene #12259), and the VSVG envelope glycoprotein vector pMD2-G (Addgene plasmid #12259) into HEK293T cells, using calcium phosphate precipitation. Supernatants of culture media were collected 48-h after transfection and filtered through a 0.45 µm syringe filter. Viral particles were then stored at −80°C until use. Efficiency of shRNA was assessed by qPCR and or western blot in case of EHMT1 and EHMT2(25). Cortical neurons were plated in a 6 well and infected at DIV 2, cell lysed and RNA extracted at DIV21.

### Reverse transcription quantitative polymerase chain reaction

RNA was extracted using the NucleoSpin^®^ RNA kit (Macherey-Nagel cat. no. 740955.50) according to the manufacturer’s protocol. Reverse transcription was performed using iScript™ cDNA Synthesis Kit (Biorad cat.no. 1708891). qPCR and melting curve analyses were performed using GoTaq^®^ qPCR Master Mix (Promega A600) on a 7500 Real-Time PCR System (Life Technologies). The primers used for qPCR are listed in Supplementary Table 7.

### Primary neuronal cell culture

One day before plating, glass coverslips (14 mm; Menzel GmbH) treated with 65% nitric acid (Sigma-Aldrich) were coated with 0.0125% polyethyleminine (PEI; Sigma-Aldrich) overnight. On the day of plating, MEAs were coated with 100 µL poly-D-lysine solution (Sigma-Aldrich, Cat.No. P7280; 100 µg/ml in 50 mM borate buffer, pH 8.5) placed directly over the array in a drop for 3 hr at 37°C in a humidified cell culture incubator. Then, MEAs were washed 3X with sterile water and air dried in a laminar flow hood. Briefly, each pregnant Wistar rat was deeply anesthetized using isoflurane (Pharmachemie B.V. Haarlem, Cat.No. 45112106) and sacrificed by cervical dislocation. Embryos (E18) were quickly and aseptically removed by caesarean section. Whole brains were removed from 2 embryos, the meninges were stripped away, and the cerebral cortices were dissected out as described(62). The tissue was placed directly into a 15 ml conical tube and digested in 2 ml papain solution (prepared as 200 U papain [Sigma-Aldrich, Cat.No. P4762], 1.6 mg L-cysteine [Sigma-Aldrich, Cat.No. C7880] in 10 mL Segal’s medium^15^) at 37°C for 45 min in a water bath. The digestion was then inactivated by adding 8 ml room-temp seeding medium (Neurobasal medium [Invitrogen, Cat.No. 21103] supplemented with 10% FBS [Sigma-Aldrich, Cat.No. F7524], 2% B-27 [Invitrogen, Cat.No. 17504], 1% GlutaMAX [Invitrogen, Cat.No. 35050] and 1% Pen/Strep solution [Sigma-Aldrich, Cat.No. P4333]). The cortices were dissociated and filtered through a 70-µm cell strainer placed atop a 50 ml conical tube. The collected cells were spun at 200 × *g* for 8 min, the medium was aspirated, and the cell pellet was gently resuspended using a pipettor with 1-ml tip and 800 µl seeding medium. The suspension was then diluted to 10 ml with seeding medium and mixed thoroughly before counting. Viable cells were counted using a hemacytometer and trypan blue exclusion. We consistently obtained 2-3 ×10^6^ cells/ml and >95% viability at the time of plating. Dissociated cells were plated on the PEI-coated glass coverslips at a final density of 1200 cells/mm^2^. For the MEAs, cells were carefully seeded onto the center of the array in a 50 µl droplet at a concentration of 1200 cells/µl. After settling for 3 hr in an incubator, 450 µL prewarmed seeding medium was slowly added to the well, followed by 500 µL culture medium (i.e. seeding medium without serum). At DIV2, cells were transduced with a lentivirus expressing GFP or an shRNA targeting the mRNA of interest. Non-infected and GFP-only infected cells were used as controls. Since we did not observe differences between these conditions, results from both were pooled together. Beginning at DIV3, and every 2 days thereafter, half the medium was removed and replenished with freshly-prepared, prewarmed culture medium.

### RNA-Seq

RNA-Seq using total RNA was executed as described in(63). In summary, 500 ng of total RNA was used to obtain double-stranded cDNA (ds-cDNA) that was subsequently purified with MinElute Reaction Cleanup Kit (Qiagen #28206). For the library construction, the KAPA Hyper Prep Kit (Kapa Biosystems #KK8504) standard protocol with a modification of a 15-minute USER enzyme (Biolab # M5505L) incubation step for the library amplification and 3ng ds-cDNA as the starting material were used. The library quantification was performed with the KAPA Library Quantification Kit (Kapa Biosystems #KK4844) followed by pair-ended sequencing with NextSeq500 (Ilumina).

### RNA-Seq data analysis

Reads were aligned using the STAR version 2.5.2b(64) to the RNOR 6.0.88 Rat Genome. UCSC genome browser tracks were obtained by generating bigwig files (using wigToBigWig) and uploading them to the UCSC genome browser(65). Count files (Read counts per gene) were then used in differential expression analysis performed with the DESeq2(35) package. Differential gene expression of KSS knockdown was calculated in comparison to GFP control knockdown. If not otherwise stated, all data analyses were performed with an adjusted P value below 0.1 (p < 0.1) to define differentially expressed genes. Principle component analysis (PCA) plots were created with the DESeq2 package. GOTERM function gene annotation analyses were performed with DAVID(66). Scatterplots were made with all genes (FPKM < 1000) and the ggplot library (http://ggplot2.org/). Venn overlap diagram was created using the Venny (version 2.1) online tool (http://bioinfogp.cnb.csic.es/tools/venny/index.html). RNAseq data are deposited to GEO: GSE120061.

### Immunocytochemistry

Primary neurons were fixed with 4% paraformaldehyde/4% sucrose in phosphate buffered saline (PBS) for 15 min. After fixation cells were washed and permeabilized with PBS and 0.2% Triton. Cells were blocked in blocking buffer (PBS, 5% normal goat serum, 1% bovine serum albumin, 0.2% Triton) for one hour, followed by incubation with primary antibodies in blocking buffer overnight at 4°C. Cells were washed with PBS the next day and incubated with secondary antibodies for one hour at room temperature. Cells were washed with PBS again to remove antibody excess and DNA was stained using Hoechst (1:10000). Finally, cells were mounted using DAKO (DAKO, S3023) fluorescent mounting medium and stored at 4°C. Epifluorescent pictures were taken at a 63x magnification using the Zeiss Axio Imager 2 equipped with apotome. All conditions within a batch were acquired with the same settings (laser power % and exposure time) in order to compare signal intensities between different experimental conditions. Number of puncta was quantified via manual counting using ImageJ (scale 1 pixel = 0.072 µm). Primary antibodies: guinea pig anti-VGAT (1:500, Synaptic Systems 131 004); mouse anti-Gephyrin (1:500, Synaptic Systems 147 111); rabbit anti-VGLUT (1:1000, Synaptic Systems 135 302); mouse anti-PSD95 (1:50, Thermo Scientific MA1-045). Secondary antibodies: goat anti-guinea pig Alexa Fluor 647 (1:2000, Invitrogen A-21450); goat anti-rabbit Alexa Fluor 647 (1:2000, Invitrogen A-21245); goat anti-mouse Alexa Fluor 568 (1:2000, Invitrogen A-11031).

### MEA recordings

The neuronal network activity was recorded for 30 min by means of micro-electrode Arrays (MEAs), devices made up of 60 planar microelectrodes (TiN/SiN, 30 mm electrode diameter, 200 mm spaced) arranged over an 8×8 square grid (except the four electrodes at the corners), supplied by Multi Channel Systems (MCS, Reutlingen, Germany). In addition, we used the 6-wells MEAs (6 independent wells, each one with 9 recording and 1 reference embedded electrodes) to perform virus dose responses experiments. After 1200x amplification (MEA 1060, MCS), signals were sampled at 10 kHz using the MCS data acquisition card. Recordings were performed outside the incubator at temperature of 37°C and to prevent evaporation and changes of the pH medium, a constant slow flow of humidified gas (5% CO_2_, 20% O_2_, 75% N_2_) was inflated onto the MEA.

### MEA data analysis

Data analysis was performed offline using a custom software package named SpyCode^16^ developed in MATLAB© (The Mathworks, Natick, MA, USA). *Spike detection.* Spike were detected by using the Precise Timing Spike Detection algorithm (PTSD)^17^. Briefly, spike trains were built using three parameters: (1) a differential threshold set to 8 times the standard deviation of the baseline noise independently for each channel; (2) a peak lifetime period (set at 2 ms); (3) a refractory period (set at 1 ms). *Burst detection.* Burst were detected using a Burst Detection algorithm^18^. The algorithm is based on the computation of the logarithmic inter-spike interval histogram in which inter-burst activity (i.e. between bursts and/or outside bursts) and intra-burst activity (i.e. within burst) for each recording channel can be easily identified, and then, a threshold for detecting spikes belonging to the same burst is automatically defined. From the spike and burst detection, the number of active channels (above threshold 0.1 spikes/sec) and the number of bursting channels (above threshold 0.4 burst/min and at least 5 spikes in burst with a minimal inter-spike-interval of 100 milliseconds) were determined. Furthermore, the mean firing rate (MFR) of the network was obtained by computing the firing rate of each channel averaged among all the active electrodes of the MEA. *Network burst detection.* Synchronous event were detected looking for sequences of closely-spaced single-channels bursts. A network burst is identified if it involves at least the 80% of the network active channels. The distributions of the network burst duration (NBD, s) and Network Inter Burst Interval (NIBI, interval between two consecutive network burst, s) were computed using bins of 10 ms and 1 s respectively. *Network burst irregularity.* Irregularity was estimated by computing the coefficient of variation (CV) of the network inter burst intervals (NIBI), which is the standard deviation divided by the mean of the NIBI.

### Whole patch clamp recordings in neuronal cultures

Experiments were performed in a recording chamber on the stage of an Olympus BX51WI upright microscope (Olympus Life Science, PA, USA) equipped with infrared differential interference contrast optics, an Olympus LUMPlan FLN 40× water-immersion objective (Olympus Life Science, PA, USA) and kappa MXC 200 camera system (Kappa optronics GmbH, Gleichen, Germany) for visualisation. Through the recording chamber a continuous flow of carbogenated artificial cerebrospinal fluid (ACSF) containing (in mM) 124 NaCl, 1.25 NaH_2_PO_4_, 3 KCl, 26 NaHCO_3_, 11 Glucose, 2 CaCl_2_, 1 MgCl_2_ (adjusted to pH 7.4), warmed to 30°C, was present. Patch pipettes (6-8 MΩ) were pulled from borosilicate glass with filament and fire-polished ends (Science Products GmbH, Hofheim, Germany) using the PMP-102 micropipette puller (MicroData Instrument, NJ, USA). Pipettes were filled with a potassium-based solution containing (in mM) 130 K-Gluconate, 5 KCl, 10 HEPES, 2.5 MgCl_2_, 4 Na_2_-ATP, 0.4 Na_3_-ATP, 10 Na-phosphocreatine, 0.6 EGTA (adjusted to pH 7.25 and osmolarity 304 mOsmol) for recordings of intrinsic electrophysiological properties in current clamp mode. For recordings of miniature inhibitory postsynaptic currents (mIPSCs) in voltage clamp mode pipettes were filled with a cesium-based solution containing (in mM) 115 CsMeSO_3_, 20 CsCl, 10 HEPES, 2.5 MgCl_2_, 4 Na_2_-ATP, 0.4 Na_3_-ATP, 10 Na-phosphocreatine, 0.6 EGTA (adjusted to pH 7.2 and osmolarity 304 mOsmol). Recordings were made using a SEC-05X amplifier (NPI Electronic GmbH, Tamm, Germany) and recorded with the data acquisition software Signal (CED, Cambridge, UK). Recordings were not corrected for a liquid junction potential of approximately −10 mV. Intrinsic electrophysiological properties were analysed using Signal and MatLab (MathWorks, MA, USA), while mIPSCs were analysed using MiniAnalysis (Synaptosoft Inc, Decatur, GA, USA).

Intrinsic electrophysiological properties were recorded in current clamp mode, where the resting membrane potential (V_rmp_) was determined after achieving whole-cell configuration. Cells were selected on a V_rmp_ of −55 mV and lower. All further recordings were performed at a holding potential of −60 mV. Passive membrane properties were determined via a 0.5 s hyperpolarizing current of −25pA. Action potential (AP) characteristics were determined from the first AP elicited by a 0.5 s depolarising current injection just sufficient to reach AP threshold. The mIPSCs were measured in voltage clamp mode, in the presence of 1 µM tetrodotoxin (TTX; Tocris, Bristol, UK), 5 µM 6-cyano-7-nitroquinoxaline-2,3-dione (CNQX) and 100 µM 2-amino-5-phosphonovaleric acid (AP-V) at a holding potential of +10 mV.

### Acute slice electrophysiology

Mice of both genders were taken at ages indicated in the text ± 1 day P7 (P6-8), P14 (P13-15) or P21 (P20-22) and anesthetized using isoflurane before decapitation. Ventral slices (350 μm) were cut using a HM650V vibratome in ice cold artificial cerebrospinal fluid (ACSF) containing (in mM): 87 NaCl; 11 Glucose; 75 Sucrose; 2.5 KCl; 1.25 NaH_2_PO_4_; 0.5 CaCl_2_; 7 MgCl_2_; 26 NaHCO_3_, continuously oxygenated with 95% O_2_/5% CO_2_ and incubated for 1h at 32°C after which they were allowed to cool down to room temperature. Before recording, slices were transferred to the recording setup and incubated in recording (ACSF) containing (unless otherwise stated) (in mM): 124 NaCl, 1.25 NaH_2_PO_4_, 3 KCl, 26 NaHCO_3_, 10 Glucose, 2 CaCl_2_, 1 MgCl_2_ and continuously oxygenated with 95% O2/5% CO2 at 30°C and incubated 15 minutes prior to recording. Cells were visualized with an upright microscope (Olympus). Patch pipettes (3-5 MΩ) were made from borosilicate glass capillaries and filled with intracellular solution containing (for voltage clamp, in mM): 115 CsMeSO_3_; 20 CsCl; 10 HEPES; 2.5 MgCl_2_; 4 Na_2_ATP; 0.4 NaGTP; 10 Na-Phosphocreatine; 0.6 EGTA (pH 7.2-7.3, 290 mOsm) or (for current clamp, in mM): 130 K-Gluconate; 5 KCl; 10 HEPES; 2.5 MgCl_2_; 0.6 EGTA; 4 Na_2_ATP; 0.4 Na_3_GTP; 10 Na-phosphocreatine (pH 7.2-7.3, 290 mOsm). Traces were recorded using a Multiclamp 700B amplifier (Molecular Devices, Wokingham, United Kingdom), sampled at 20 kHz and filtered at 2 kHz. Cells were excluded from analysis if the access resistance exceeded 25 MΩ. Miniature inhibitory postsynaptic currents (mIPSCs) were recorded in the presence of Tetrodotoxin (1 µM), 6-Cyano-7-nitroquinoxaline-2,3-dione (CNQX, 5 µM) and D-(-)-2-Amino-5-phosphonopentanoic acid (D-APV 100 µM). Miniature excitatory postsynaptic currents (mEPSCs) were recorded in the presence of TTX and Picrotoxin (PTX, 100 µM). Paired pulse ratio (PPR) was recorded in the presence of CNQX and D-APV, the recording ACSF contained 4 mM CaCl_2_ and 4 mM MgCl_2_, and was calculated as peak2/peak1 after correcting for any residual current at the second pulse. Spontaneous action potential (AP) frequency was calculated from the total number of APs during a 10 minute recording. Intrinsic properties were analyzed as follows: Resting membrane potential was recorded directly after break in. All other properties were recorded at a holding potential of −60 mV. Input resistance was calculated from 6 responses to increasing negative current injections (5 pA per step)(67). Rise and decay time were calculated at 20%-80%/80%-20% of the amplitude, respectively. The adaptation ratio was calculated as the 8th/3rd inter-spike interval. Miniature recordings were analyzed using Mini Analysis Program (Synaptosoft, Decatur, GA, USA). Other traces were analyzed using Clampfit 10.2. All drugs were purchased from Tocris (Abingdon, United Kingdom).

### Animals

For the experiments presented in this study, mice heterozygous for a targeted loss-of-function mutation in the *Ehmt1* gene (*Ehmt1+/-* mice) and their WT littermates on C57BL/6 background were used, as previously described(20). Animal experiments were conducted in conformity with the Animal Care Committee of the Radboud University Nijmegen Medical Centre, The Netherlands, and conform to the guidelines of the Dutch Council for Animal Care and the European Communities Council Directive 2010/63/EU.

### Statistics

The statistical analysis of the MEA data were performed using OriginPro (OriginLab Corporation, Northampton, MA, USA) and MATLAB© (The Mathworks, Natick, MA, USA). Statistical analysis of the single cells data were performed using GraphPad Prism 5 (GraphPad Software, Inc., CA, USA). We determined normal distribution using a Kolmogorov-Smirnov normality test. Depending on the distribution of the results, we determined the statistical significance for the different experimental conditions by either a parametric (paired Student’s t-test) or non-parametric (Kruskal-Wallis or Mann-Whitney test) tests where p-value < 0.05 were considered significant. Significance over development was determined with a one-way ANOVA when normal Gaussian distribution was observed or Kruskal-Wallis test when this was not applicable. Data are expressed as Mean ± standard error of the mean (SE).

### Study approval

Animal experiments were conducted in conformity with the Animal Care Committee of the Radboud University Nijmegen Medical Centre, The Netherlands, and conform to the guidelines of the Dutch Council for Animal Care and the European Communities Council Directive 2010/63/EU.

## Author contributions

D.S, H.v.B and N.N.K conceived and supervised the study. M.S performed all animal work. B.M, M.F, M.S, A.O, J.K, E.B and NNK designed and performed all experiments. T.K. provided resources. H.Z, J.Q, P.K, R.M, B.M, M.F and S.J. performed data analysis. D.S, M.F, B.M, M.S, H.v.B, T.K, H.Z and N.N.K wrote the manuscript.

## Acknowledgements

This work was supported by grants from of the Netherlands Organization for Scientific Research (open ALW ALW2PJ/13082 to H.v.B. and N.N.K.) and the Jerome Lejeune Foundation (to H.v.B).

---

**Supplementary Figure S1.**
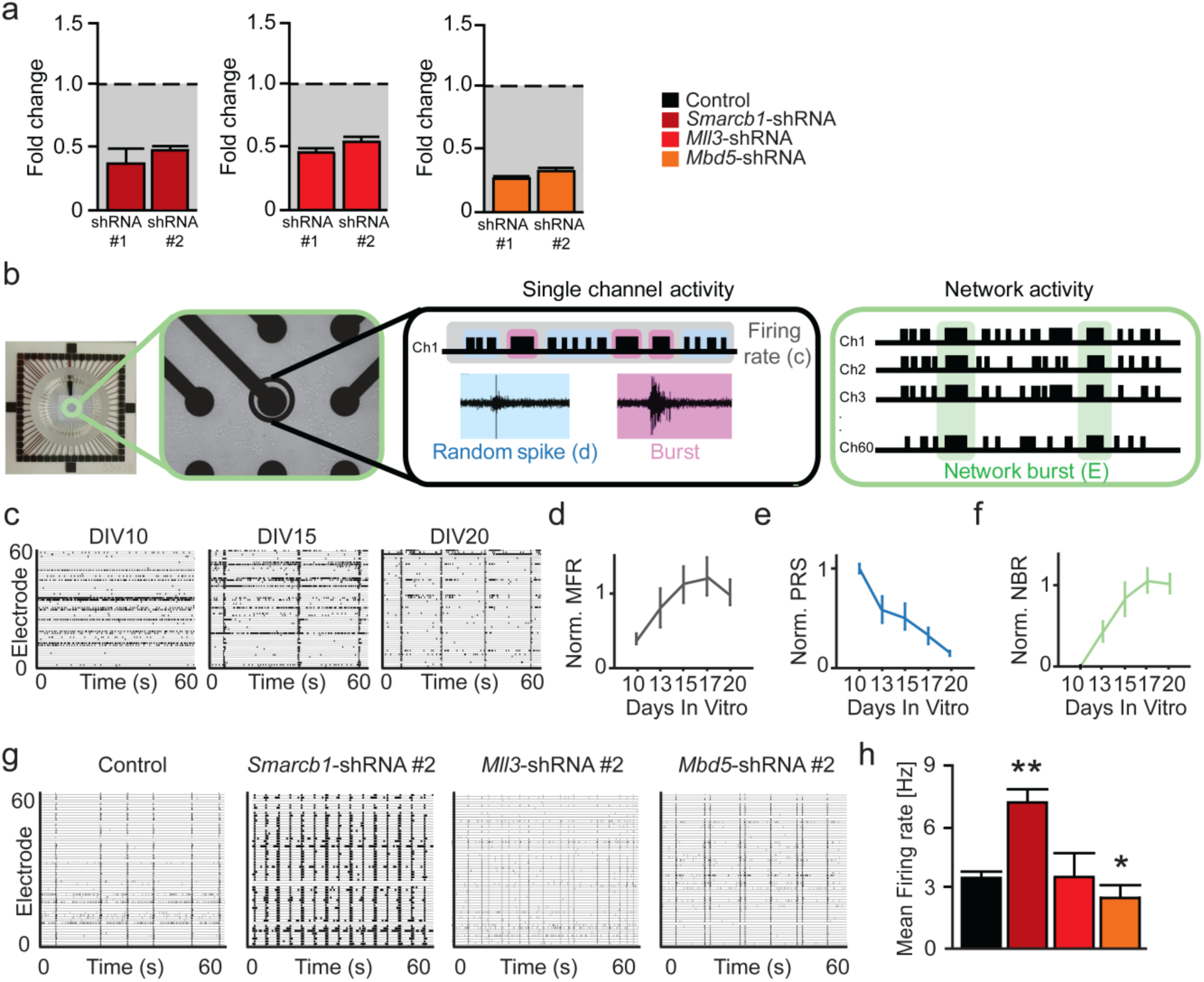
Characterization of secondary hairpins. **a.** Quantitative PCR of KSS genes on cortical primary neurons at DIV21. Cells were infected at DIV2 with respective lentivirus expressing shRNA. Data show a reduction of mRNA levels for both hairpins targeting one of KSS genes. **b.** Schematic overview of an electrophysiological recording from cells cultured on Micro-electrode arrays (MEAs). Neuronal activity is recorded through the embedded microelectrodes (first green box). From one single electrode (single channel activity, black box), either spikes or burst can be detected. From all events in this channel, the mean firing rate (MFR, c) and the percentage random spikes (PRS, d) can be calculated. From all electrodes in the MEA (second green box), synchronous network activity can be detected (network burst). From all synchronous events, the network burst rate (NBR, e) can be calculated. **c.** Representative raster plots showing 60 s of spontaneous activity exhibited by control cultures at different time points. **d-f.** Control neuronal network development normalized to values of DIV17. **d.** Normalized Mean Firing Rate (MFR) of control networks shows increasing amounts of total events during development reaching a plateau at DIV 17 (n=10). **e.** Normalized Percentage of Random Spikes (PRS) of control networks show decreasing amount of random spiking activity over development (n=10). **f.** Normalized Network Burst Rate (NBR) of control networks show increasing amounts of network bursts over development reaching a plateau at DIV 17 (n=10). **g.** Representative raster plots showing 60 s of electrophysiological activity exhibited by control- and KSS gene*–* deficient cultures (second independent shRNA) at DIV 20. **h.** Graph showing the mean firing rate for KSS gene-deficient cultures at DIV 20. The mean firing rate is significantly increased in SMARCB1-deficient cultures and decreased in MBD5-deficient cultures. No significant change was found for MLL3-deficient cultures (n=16 for controls, n=4 for *Smarcb1*-shRNA, n=3 for *Mll3*-shRNA and n=5 for *Mbd5*-shRNA). Sample size (n) indicates number of MEA experiments. Data represents mean ± SEM .* p < 0.05, ** p < 0.01 (parametric two-tailed T-test was performed between controls and KSS gene-deficient cultures).

**Figure S2.**
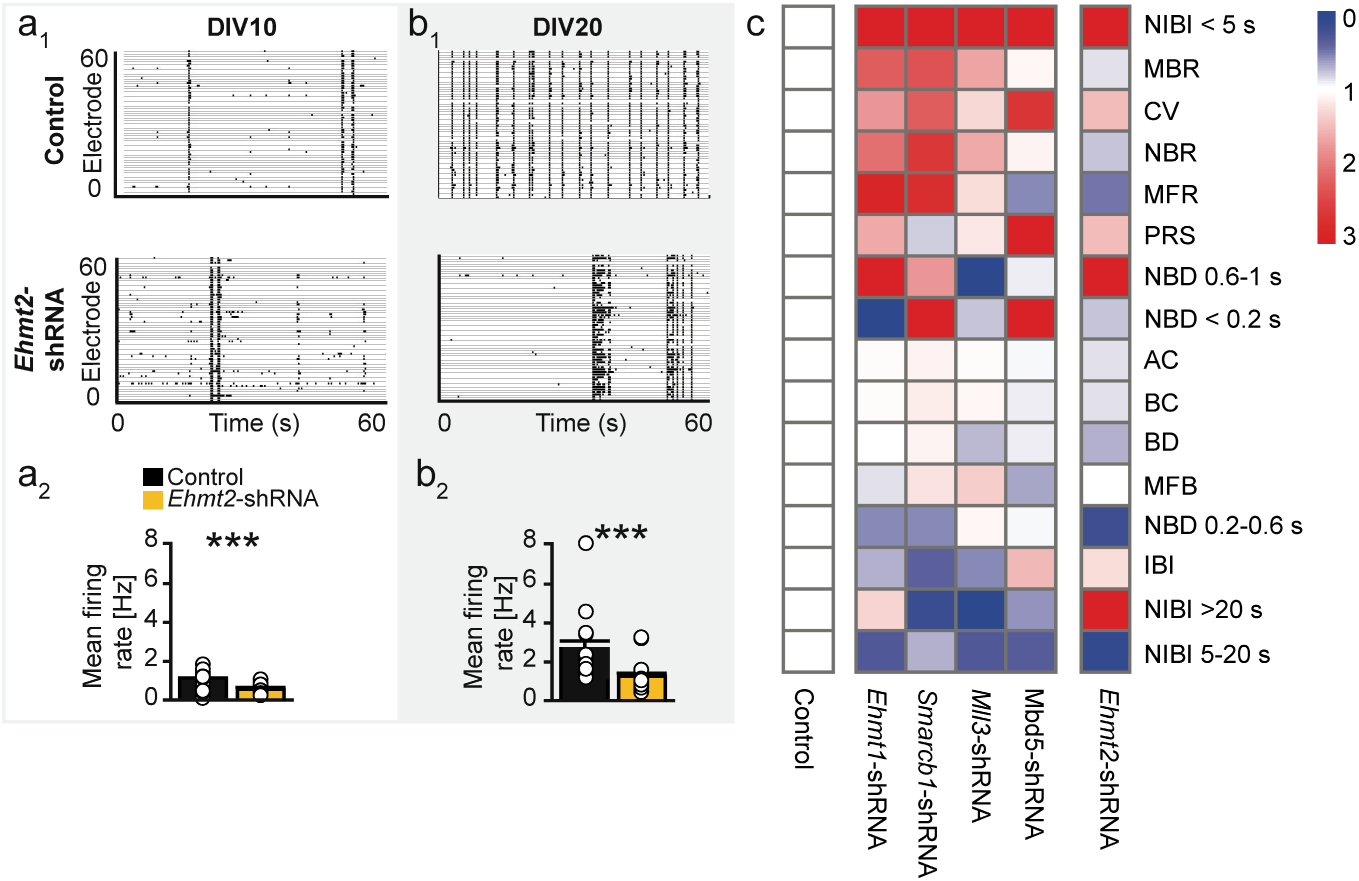
EHMT2-deficient cultures show distinct phenotypic differences over development. **a1.** Representative raster plots from control cultures and EHMT2-deficient cultures at DIV10, displaying 60 seconds of spontaneous activity (x-axis) for each individual electrode (i.e. 60 electrodes on y-axis). **a2**. Mean firing rate is significantly decreased in EHMT2-deficient cultures at DIV10 as compared to controls. **b1.** Representative raster plots from control- and EHMT2-deficient cultures at DIV 20. **b2**. Mean firing rate is significantly decreased in EHMT2-deficient cultures at DIV20. **c.** Heatmap showing the relative values of the parameters describing the phenotype exhibited by EHMT1-, SMARCB1-, MLL3-, MBD5- and EHMT2-deficient neuronal network as compared to control. The scale of the relative values is indicated from 0 (blue) to 3 (red), where 1 indicates the control (white) (n=16 for controls and n=15 for EHMT2-deficient cultures). Sample size (n) indicates the number of cells per genotype or the number of recorded MEA experiments over development. DIV: Days in Vitro. Data represents mean ± SE. *** p < 0.001 (either a nonparametric Kruskal Wallis or a parametric two-tailed T-test was performed between KSS gene-deficient cultures and matched controls).

**Figure S3.**
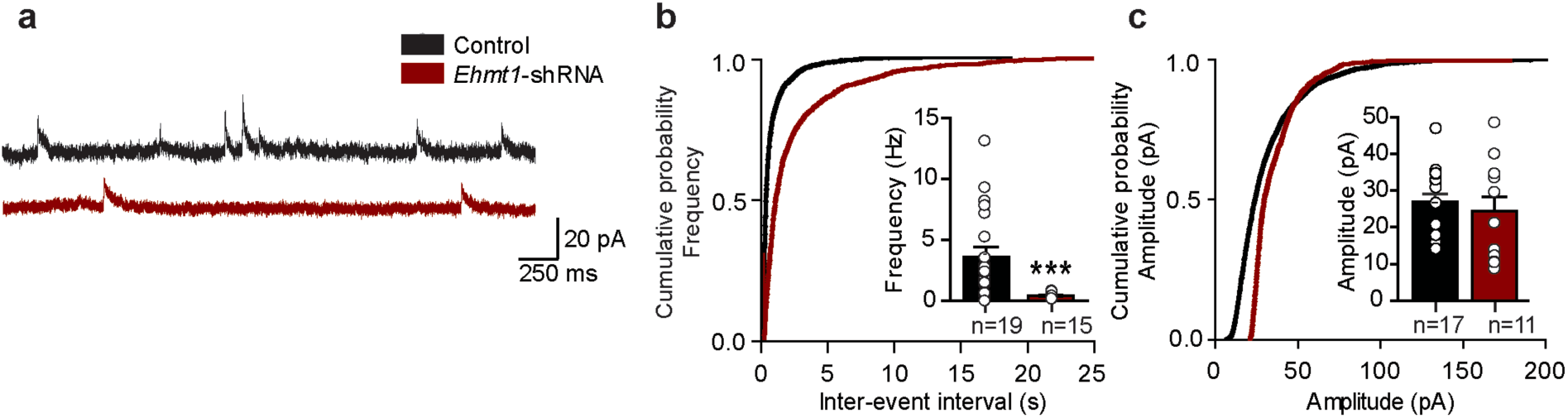
Reduced inhibitory input in EHMT1-deficient rat cortical networks. **a**. Example traces of mIPSC recordings from EHMT1-deficient and control cultures at DIV20. **b**. Quantification of mIPSC frequency from EHMT1-deficient (n=19) and control cultures (n=15) at DIV20. **c**. Quantification of mIPSC amplitude from EHMT1-deficient (n=17) and control (n=11) cultures at DIV20. Sample size (n) is indicated in the bars as number of cells. Data represents mean ± SE. *** p < 0.001. (either a nonparametric Kruskal Wallis or a parametric two-tailed T-test was performed between KSS gene-deficient cultures and matched controls).

**Figure S4.**
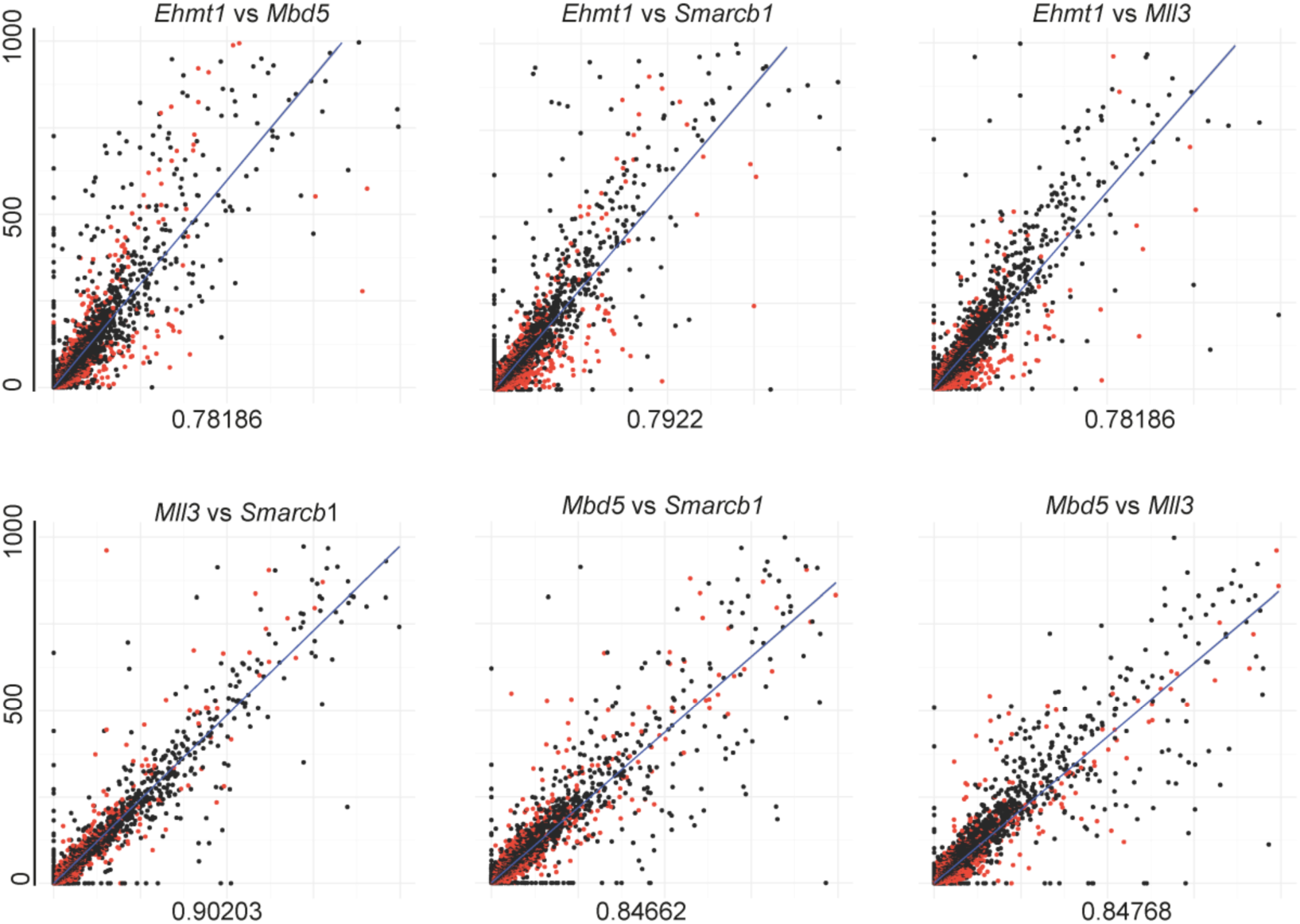
Scatter plots of all pair-wise comparisons between samples. R2 adjusted values are given below the plots. The label above each plot shows which two samples are depicted, with the 1^st^ sample represented by the Y axis and the 2^nd^ represented by the X axis. The scatter plots were made using raw FPKM values of all genes (FPKM < 1000). In black are non-differentially expressed genes (NDE) and in orange differentially expressed (DE) ones.

**Figure S5.**
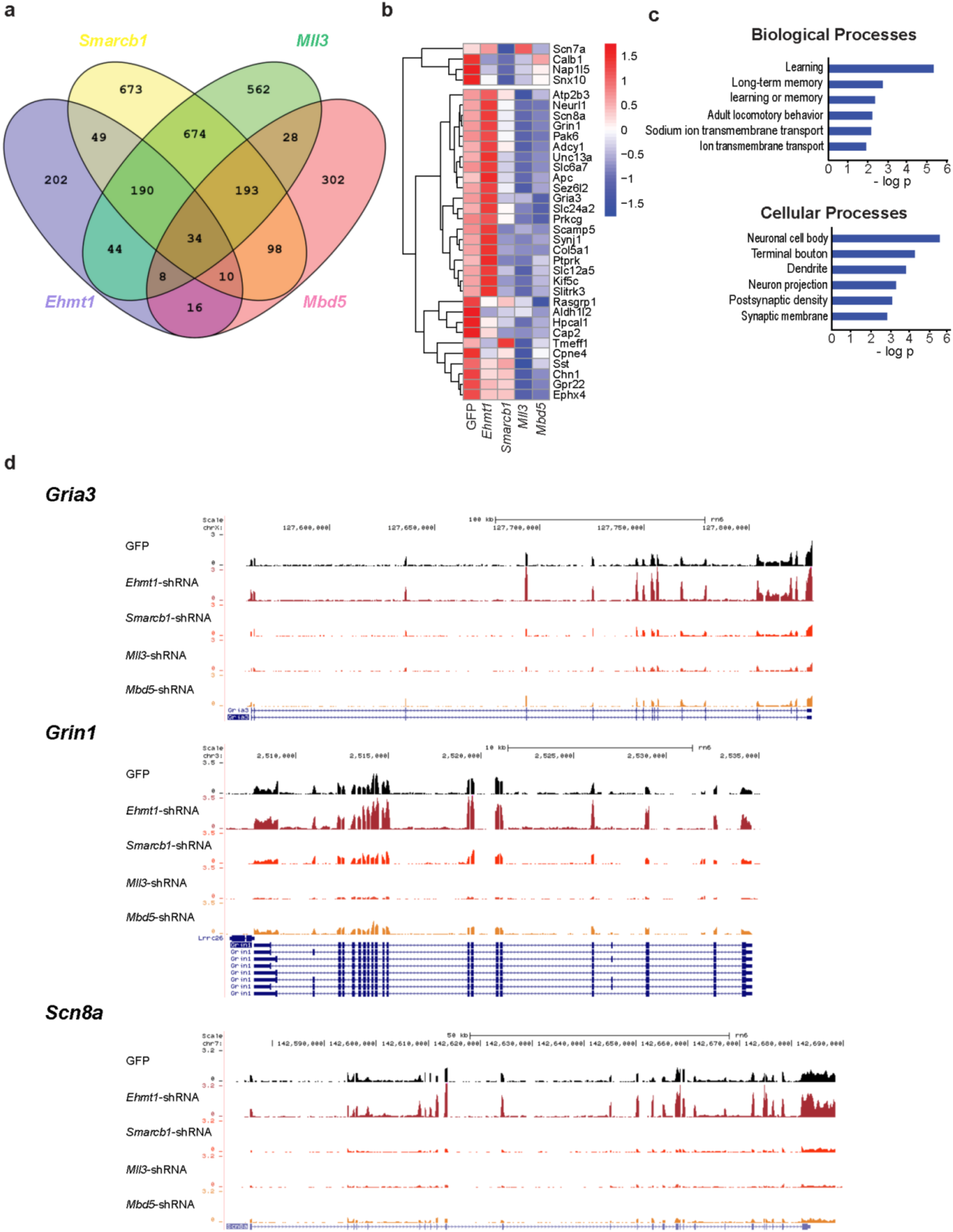
Overlap analysis shows 34 overlapping DE genes detected in the four knock-down samples, as compared to the control (GFP). **a.** Venn diagram of DE genes (q < 0.1) from the four knock-down samples and their overlaps. **b.** Heatmap of the relative expression (Z scores) of the 34 overlapping DE genes detected in the four knockdown samples, as compared to the control (GFP). The scale of the Z scores is indicated from −1.5 (blue) to 1.5 (red). **c.** Top 6 GO terms detected from the 34 DE genes in the categories of ‘biological processes’ and ‘cellular processes’. **d.** UCSC genome browser screenshots showing the RNA-seq signals at the loci of *Gria3, Grin1* and *Scn8A* in all samples.

**Figure S6.**
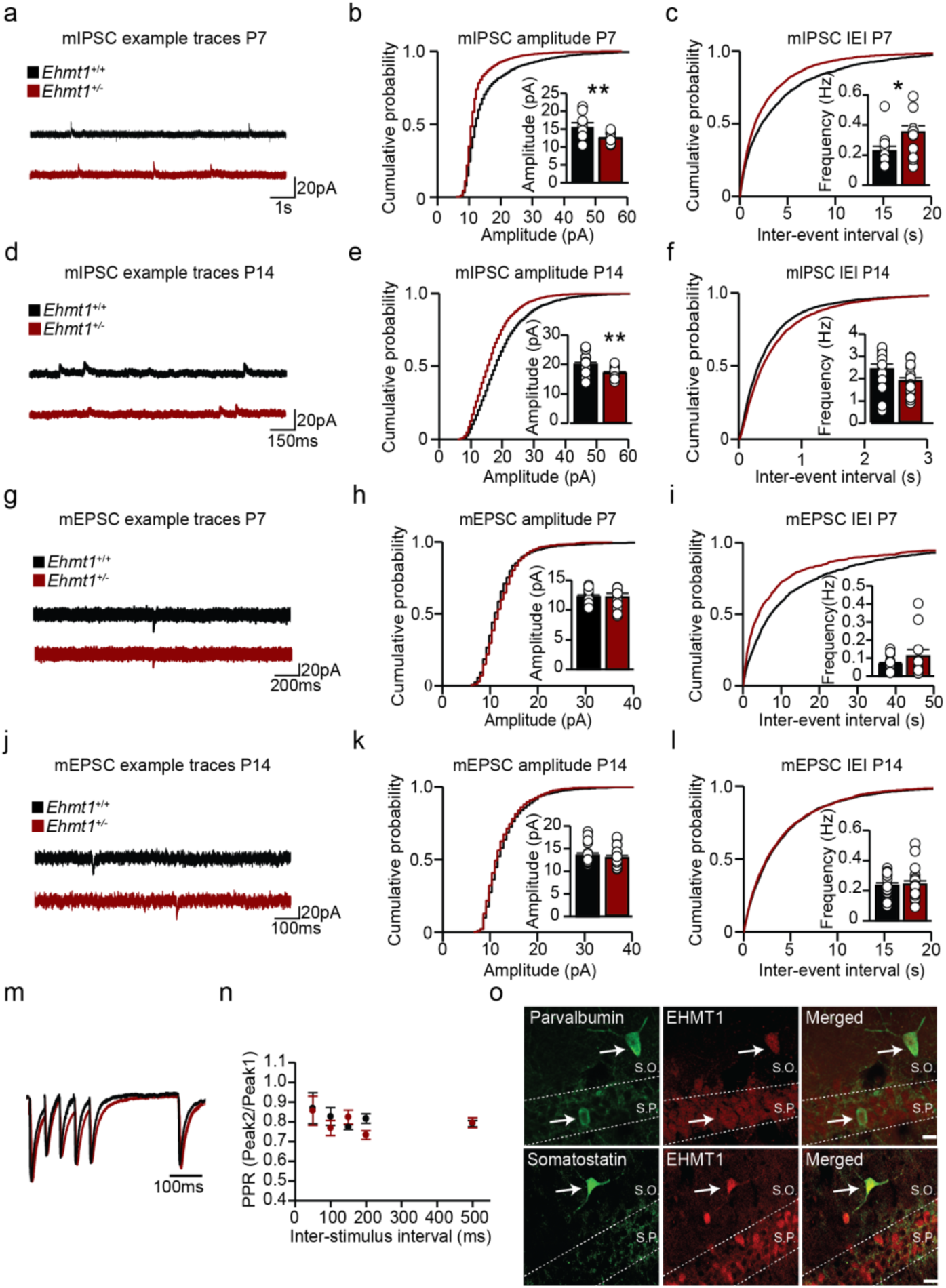
Development of synaptic connectivity of CA1 pyramidal neurons. **a.** Example traces of mIPSC recordings from *Ehmt1^+/+^* and *Ehmt1^+/-^* mice at P7. **b-c.** Quantification of mIPSC amplitude (**b**) and frequency (**c**) from *Ehmt1^+/+^* (n=11) and *Ehmt1^+/-^* (n=16) mice at P7. **d.** Example traces of mIPSC recordings from *Ehmt1^+/+^* and *Ehmt1^+/-^* mice at P14. **e-f**. Quantification of mIPSC amplitude (**e**) and frequency (**f**) from *Ehmt1^+/+^* (n=17) and *Ehmt1^+/-^* (n=17) mice at P14. **g.** Example traces of mEPSC recordings from *Ehmt1^+/+^* and *Ehmt1^+/-^* mice at P7. **h-i**. Quantification of mEPSC amplitude (**h**) and frequency (**i**) from *Ehmt1^+/+^* (n=16) and *Ehmt1^+/-^* (n=11) mice at P7. **j.** Example traces of mEPSC recordings from *Ehmt1^+/+^* and *Ehmt1^+/-^* mice at P14. **k-l**. Quantification of mEPSC amplitude (**k**) and frequency (**l**) from *Ehmt1^+/+^* (n=18) and *Ehmt1^+/-^* (n=19) mice at P14. **m**. Example traces of PPR responses following stimulation of the SO at P21. **n**. Quantification of PPR responses following stimulation in SO at P21. **o**. Immunohistochemical analysis showing colocalization of EHMT1 with both PV and SOM in SO and SP. Scale bar indicated 10 μm. SO: Stratum oriens. SP: Stratum pyramidale. IEI: Inter-event interval. PPR: Paired-pulse ratio. Sample size (n) indicates the number of cells. * p < 0.05; ** p < 0.01. Data represents mean ± SE, Two-tailed Student’s t-test.

**Supplementary Table 1.** Values from **Figure 1**. Data represents mean ± SE. Either a nonparametric Kruskal Wallis or a parametric two-tailed T-test was performed between KSS gene-deficient cultures and matched controls.

| Fig a |  | DIV | Control | *Ehmt1*-shRNA | P-value |
| --- | --- | --- | --- | --- | --- |
| 2 | MFR (Hz) | 10 | 1,2±0,16 | 1,63±0,16 | 0,082 |
| 3 | PRS (%) | 10 | 71,57±3,23 | 87,94±3,91 | 0,022 |
| 4 | NBR (NB/min) | 10 | 0,01±0,01 | 0,4±0,2 | 0,103 |
| 5 | MFR (Hz) | 20 | 2,42±0,34 | 6,97±1,58 | 0,018 |
| 6 | PRS (%) | 20 | 11,01±2,41 | 17,66±2,56 | 0,077 |
| 7 | NBR (NB/min) | 20 | 3,62±0,41 | 7,40±1,44 | 0,031 |
| Fig b |  | DIV | Control | *Smardc1*-shRNA | P-value |
| 2 | MFR (Hz) | 10 | 1,2±0,16 | 1,22±0,19 | 0,93 |
| 3 | PRS (%) | 10 | 71,57±3,23 | 92,14±1,38 | 0,0004 |
| 4 | NBR (NB/min) | 10 | 0,01±0,01 | 0,23±0,13 | 0,19 |
| 5 | MFR (Hz) | 20 | 2,42±0,34 | 6,57±0,89 | 0,0003 |
| 6 | PRS (%) | 20 | 11,01±2,41 | 8,90±1,28 | 0,46 |
| 7 | NBR (NB/min) | 20 | 3,62±0,41 | 9,37±1,66 | 0,0025 |
| Fig c |  | DIV | Control | *Mll3*-shRNA | P-value |
| 2 | MFR (Hz) | 10 | 1,35±0,17 | 1,44±0,14 | 0,72 |
| 3 | PRS (%) | 10 | 34,46±5,44 | 53,23±5,75 | 0,048 |
| 4 | NBR (NB/min) | 10 | 1,73±0,31 | 1,18±0,12 | 0,21 |
| 5 | MFR (Hz) | 20 | 6,32±1,10 | 7,86±0,74 | 0,25 |
| 6 | PRS (%) | 20 | 4,97±0,70 | 5,94±1,13 | 1,000 |
| 7 | NBR (NB/min) | 20 | 6,97±0,79 | 11,33±0,96 | 0,006 |
| Fig d |  | DIV | Control | *Mbd5*-shRNA | P-value |
| 2 | MFR (Hz) | 10 | 1,35±0,17 | 2,71±0,25 | 0,001 |
| 3 | PRS (%) | 10 | 34,46±5,44 | 79,06±2,11 | 1,65E-07 |
| 4 | NBR (NB/min) | 10 | 1,73±0,31 | 2,7±0,68 | 0,24 |
| 5 | MFR (Hz) | 20 | 6,32±1,10 | 3,64±0,6 | 0,04 |
| 6 | PRS (%) | 20 | 4,97±0,70 | 14,81±1,87 | 2,00E-04 |
| 7 | NBR (NB/min) | 20 | 6,97±0,79 | 7,79±1,66 | 0,67 |

**Supplementary Table 2.** Values from **Figure 3**. Data represents mean ± SE. Either a nonparametric Kruskal Wallis or a parametric two-tailed T-test was performed between KSS gene-deficient cultures and matched controls.

|  |  | Control | *Ehmt1*-<br>shRNA | P-value | *Smarchb1*-<br>shRNA | P-value | *Mll3*-<br>shRNA | P-value | *Mbd5*-<br>shRNA | P-value |
| --- | --- | --- | --- | --- | --- | --- | --- | --- | --- | --- |
| **b** | RMP (mV) | -61,5±0,6 | -61,2±1,0 | 0,79 | -57,9±0,9 | -57,9 | -62,8±1,2 | 0,27 | -58,4±1,1 | 0,05 |
| **c** | Input resistance<br>(mΩ) | 285,3±16,7 | 226,6±17,0 | 0,02 | 310,7±28,7 | 310,7 | 368,0±26,6 | 0,01 | 376,8±31,8 | 0,047 |
| **d** | Threshold (mV) | -31,6±0,9 | -35,5±0,8 | 0,00 | -34,5±0,8 | -34,5 | -35,2±1,0 | 0,01 | -30,9±0,7 | 0,49 |
| **e** | Rheobase (pA) | 90,2±7,0 | 81,4±8,5 | 0,43 | 55,4±5,5 | 55,4 | 47,5±3,8 | 0,00 | 65,9±8,8 | 0,06 |
|  | Rise time (ms) | 0,4±0,0 | 0,4±0,0 | 0,50 | 0,4±0,0 | 0,4 | 0,5±0,0 | 0,00 | 0,5±0,0 | 0,144 |
|  | Decay time (ms) | 1,7±0,1 | 1,6±0,1 | 0,22 | 2,4±0,2 | 2,4 | 2,6±0,4 | 0,01 | 1,3±0,1 | 0,04 |
|  | Overshoot (mV) | 35,7±1,0 | 32,8±1,4 | 0,09 | 36,6±1,2 | 36,6 | 27,9±1,6 | 0,00 | 31,7±1,6 | 0,09 |
|  | Time constant (ms) | 32,5±1,6 | 26,5±1,2 | 0,01 | 32,6±2,7 | 32,6 | 33,0±2,6 | 0,98 | 38,6±3,4 | 0,21 |
|  | Amplitude (mV) | 67,2±1,2 | 68,2±1,4 | 0,59 | 71,1±1,2 | 71,1 | 63,1±1,7 | 0,05 | 62,9±1,4 | 0,05 |
|  | Capacitance (pF) | 120,8±6,5 | 124,7±8,2 | 0,71 | 107,7±5,9 | 107,7 | 91,9±6,7 | 0,01 | 109,1±7,9 | 0,30 |
|  | Duration at 50%<br>(ms) | 2,1±0,1 | 1,8±0,1 | 0,05 | 2,6±0,2 | 2,6 | 2,7±0,2 | 0,00 | 1,7±0,1 | 0,08 |
| **h** | VGAT-Gephyrin co-<br>localised<br>Puncta/10µm | 4,39±0,18 | 3,06±0,25 | <0,00<br>01 | 2,35±0,42 | <0,00<br>01 | 1,66±0,43 | <0,00<br>01 | 1,40±0,12 | <0,00<br>01 |
|  | VGLUT-PSD-95 co-<br>localised<br>Puncta/10µm | 3,86±0,16 | 3,70±0,62 | 0,934<br>2 | 2,21±0,37 | <0,00<br>01 | 1,64±0,29 | <0,00<br>01 | 1,70±0,13 | <0,00<br>01 |

**Supplementary Table 3:** List of differentially expressed genes upon KSS gene knockdown **Supplementary Table 4:** List of common dysregulated genes

**Supplementary Table 5.** Table related to **Figure 4**. Overview of 34 overlapping DE genes detected in the four knock-down samples.

| Gene | Description | Relevant associations | References |
| --- | --- | --- | --- |
| ***Adcy1*** | Adenylate cyclase 1 | ASD | 1,2 |
| ***Aldh1l2*** | Aldehyde dehydrogenase 1 family, member L2 |  |  |
| ***Apc*** | Adenomatous polyposis coli | ASD, ID | 3,4 |
| ***Atp2b3*** | ATPase plasma membrane Ca<sup>2+</sup> transporting 3; PMCA3 | sleep duration | 5 |
| ***Calb1*** | Calbindin 1 | Alzheimer disease, epilepsy | 6 |
| ***Cap2*** | Adenylyl cyclase-associated protein 2 |  |  |
| ***Chn1*** | Chimerin 1 | Duane's retraction syndrome | 7 |
| ***Col5a1*** | Collagen type V alpha 1 chain |  |  |
| ***Cpne4*** | Copine 4 |  |  |
| ***Ephx4*** | Epoxide hydrolase 4 |  |  |
| ***Gpr22*** | G protein-coupled receptor 22 |  |  |
| ***Gria3*** | Glutamate ionotropic receptor AMPA type subunit 3; GluA3 | ID, sleep disturbances | 8 |
| ***Grin1*** | Glutamate ionotropic receptor NMDA type subunit 1; GluN1 | ID, hypotonia | 9 |
| ***Hpcal1*** | Hippocalcin-like 1 | ASD | 10 |
| ***Kif5c*** | Kinesin family member 5C | ID, epilepsy, microcephaly, cortical malformation | 11,12 |
| ***Nap1l5*** | Nucleosome assembly protein 1-like 5 |  |  |
| ***Neurl1*** | Neuralized E3 ubiquitin protein ligase 1 |  |  |
| ***Pak6*** | p21 (RAC1) activated kinase 6 | neurite formation | 13,14 |
| ***Prkcg*** | Protein kinase C, gamma | ASD, spinocerebellar ataxia | 15,16,17 |
| ***Ptpnk*** | Protein tyrosine phosphatase, receptor type, K |  |  |
| ***Rasgrp1*** | RAS guanyl releasing protein 1 | Huntington disease, Parkinson disease | 18 |
| ***Scamp5*** | Secretory carrier membrane protein 5 | ASD | 19,20,21 |
| ***Scn7a*** | Sodium voltage-gated channel alpha subunit 7 | excitability in sensory neurons | 22 |
| ***Scn8a*** | Sodium voltage-gated channel alpha subunit 8; Na<sub>v</sub>1.6 | ID, epilepsy | 23 |
| ***Sez6l2*** | Seizure related 6 homolog like 2 | ASD | 24 |
| ***Slc12a5*** | Solute carrier family 12 member 5; KCC2 | ASD, Rett syndrome, schizophrenia | 25,26,27,28 |
| ***Slc24a2*** | Solute carrier family 24 member 2; NCKX2 | learning and memory, Alzheimer disease | 29 |
| ***Slc6a7*** | Solute carrier family 6 member 7 | ID | 30 |
| ***Slitrk3*** | SLIT and NTRK-like family, member 3 | schizophrenia, anxiety disorders, obsessive compulsive disorders | 31 |
| ***Snx10*** | Sorting nexin 10 | dopaminergic neurons | 32 |
| ***Sst*** | Somatostatin | ASD | 33,34 |
| ***Synj1*** | Synaptojanin 1 | ASD, epilepsy | 19,28,35 |
| ***Tmeff1*** | Transmembrane protein with EGF-like and two follistatin-like domains 1; Tomoregulin-1 | Parkinson disease; dopaminergic neurons | 36 |
| ***Unc13a*** | Unc-13 homolog A; Munc13-1 | ASD, ID, dyskinesia | 27,28,37 |

**Supplementary Table 6.** Values from **Figure 5**. Data represents mean ± SE. Either a nonparametric Kruskal Wallis or a parametric two-tailed T-test was performed between KSS gene-deficient cultures and matched controls.

| Fig |  | WT |  |  | HET |  |  | P-value |
| --- | --- | --- | --- | --- | --- | --- | --- | --- |
| **k** | RMP (mV) | -67.21 | ± | 0.76 | -66.62 | ± | 0.82 | 0.618 |
|  | Capacitance (pF) | 184.69 | ± | 9.27 | 170.39 | ± | 12.83 | 0.374 |
|  | AP Peak (mV) | 45.47 | ± | 0.90 | 46.17 | ± | 0.97 | 0.621 |
|  | Input resistance (MΩ) | 210.90 | ± | 10.02 | 191.02 | ± | 7.72 | 0.525 |
|  | Half width (ms) | 0.92 | ± | 0.02 | 0.91 | ± | 0.02 | 0.520 |
|  | Rise Time (ms) | 0.16 | ± | 0.00 | 0.15 | ± | 0.01 | 0.301 |
|  | Decay Time (ms) | 0.52 | ± | 0.01 | 0.51 | ± | 0.01 | 0.905 |
|  | AP threshold (mV) | -36.98 | ± | 0.72 | -42.64 | ± | 0.83 | 0.000 |
| **n** | Rheobase (pA) | 42.11 | ± | 2.64 | 23.85 | ± | 2.99 | 0.000 |
|  | AP adaptation ratio | 1.70 | ± | 0.07 | 1.89 | ± | 0.17 | 0.244 |

**Supplementary Table 7:**
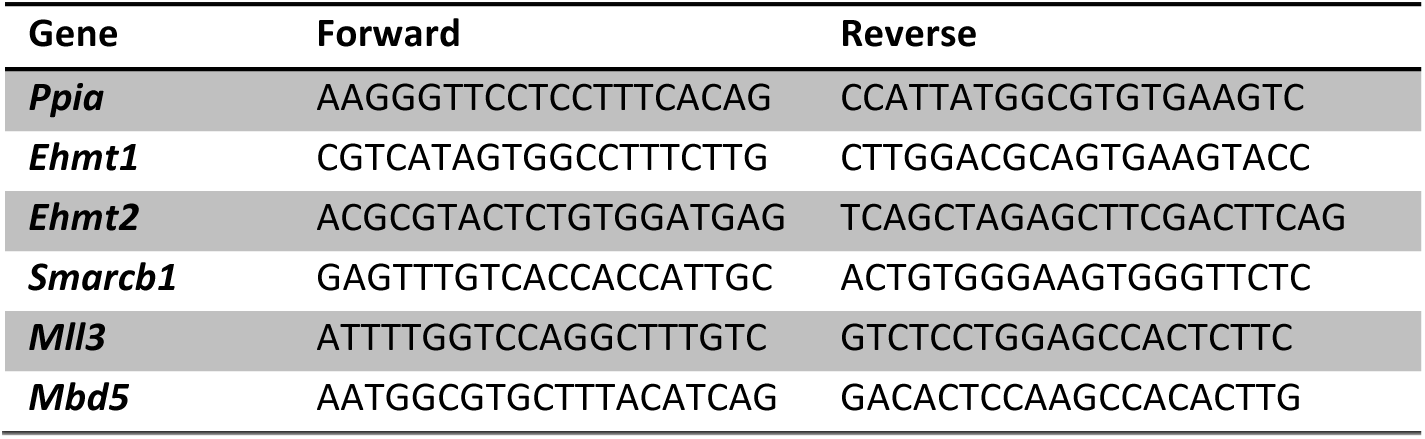
sequence information of qPCR primers.

